## Supporting Information for "Structural insights into substrate recognition by the SOCS2 E3 ubiquitin ligase"

### Supplementary Figures and Tables

(a) (b) (c)

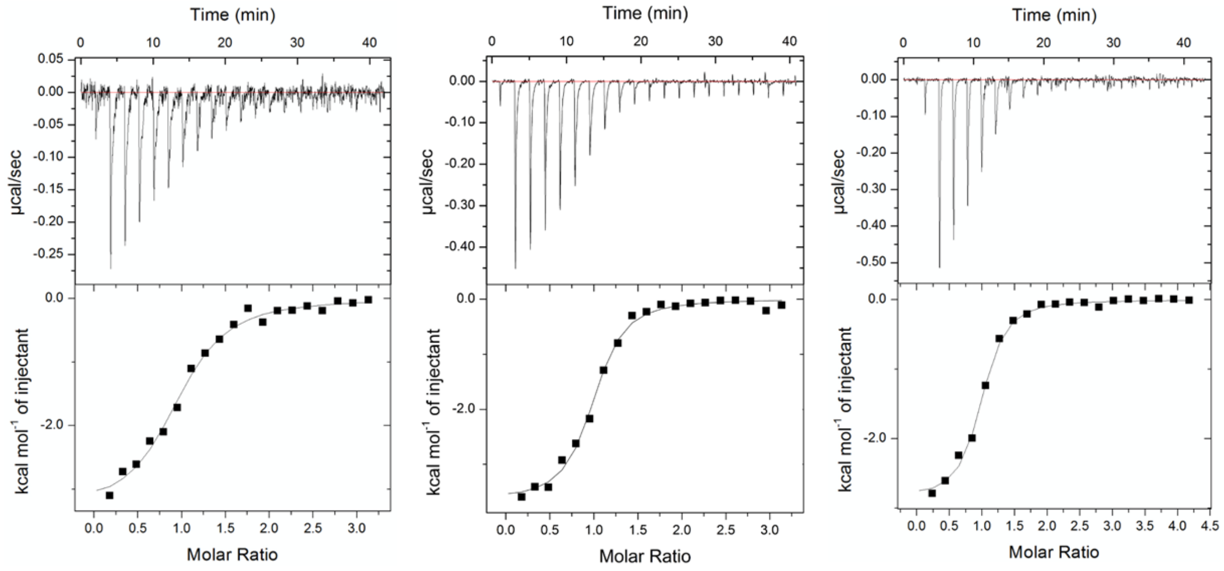


(d)

| SBC v/s | No. of experiments | $K_D$ (µM) | $\Delta G$ (kcal × mol$^{-1}$) | $\Delta H$ (kcal × mol$^{-1}$) | T$\Delta S$ (kcal × mol$^{-1}$) | Stoichiometry (N) |
|---|---|---|---|---|---|---|
| **GHR_pY595*** | **4** | **1.11 ± 0.15** | **-8.12 ± 0.08** | **-3.48 ± 0.07** | **4.63 ± 0.08** | **1.03 ± 0.03** |
| EpoR_pY426[$] | 7 | 6.92 ± 1.27 | -7.04 ± 0.11 | -3.14 ± 0.18 | 3.84 ± 0.21 | 1.11 ± 0.04 |
| **GHR_pY487[#]** | **3** | **2.30 ± 0.37** | **-7.69 ± 0.09** | **-2.82± 0.06** | **4.84± 0.11** | **0.93 ± 0.03** |

*GHR_pY595, PVPDpYTSIHIV-amide
[$]EpoR_pY426, ASFEpYTILDPS-amide
[#]GHR_pY487, NIDFpYAQVSDI -amide

**Figure S1. Biophysical characterisation of the interactions between SBC and phosphorylated substrate peptides**

ITC measurement of (a) the EpoR_pY426 peptide (b) GHR_pY595 peptide and (c) GHR_pY487 peptide binding to the SOCS2-EloB-EloC ternary (SBC) complex at 298K. (d) ITC binding data for phosphorylated substrate peptides. Values reported are the means ± s.e.m. from independent experiments.

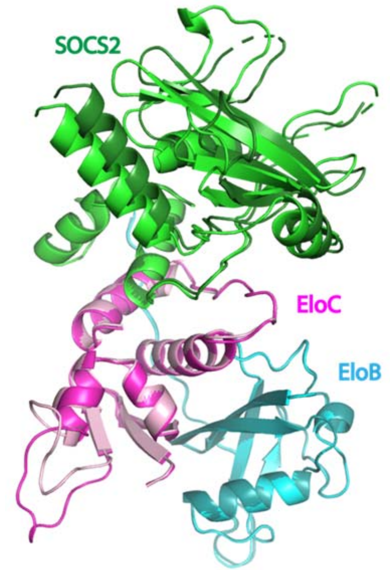


**Figure S2. The hinge motion of SOCS2**

Superposition of the two protomers from SBC-GHR structure via EloB (cyan and blue) backbone atom alignment. A hinge motion of SOCS2 (green and dark green) is observed.

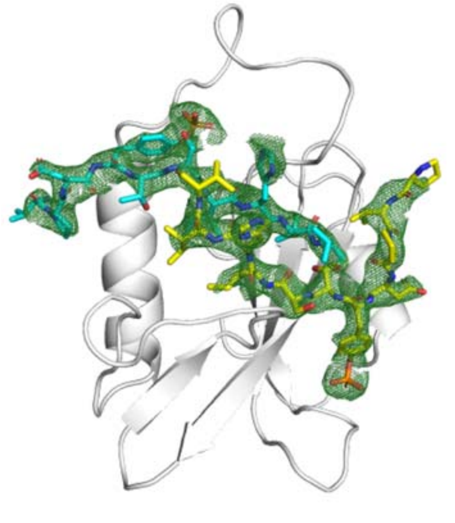

**Figure S3. Structural detail of the SBC-GHR$_2$ co-crystal**

The Fo-Fc ligand omit map of the GHR peptides (green mesh) contoured at 2.5 σ level to highlight densities for the GHR_pY595 peptides (yellow and cyan stick).

(a)

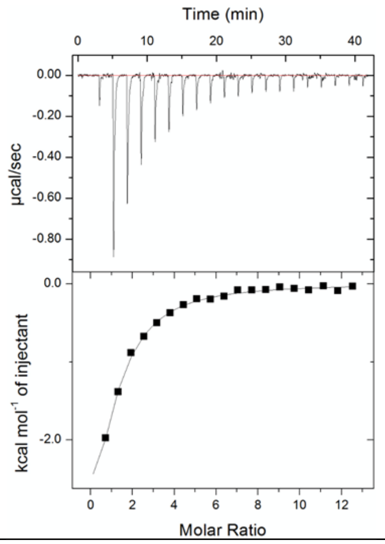


| SBC v/s | No. of experiments | $K_D$ (µM) | Δ$G$ (kcal × mol$^{-1}$) | Δ$H$ (kcal × mol$^{-1}$) | TΔ$S$ (kcal × mol$^{-1}$) | Stoichiometry (N) |
|---|---|---|---|---|---|---|
| **Spy** | 4 | 50 ± 4.44 | -5.86 ± 0.05 | -5.34 ± 0.26 | 0.52 ± 0.27 | 1.00 ± 0.09 |

(b)

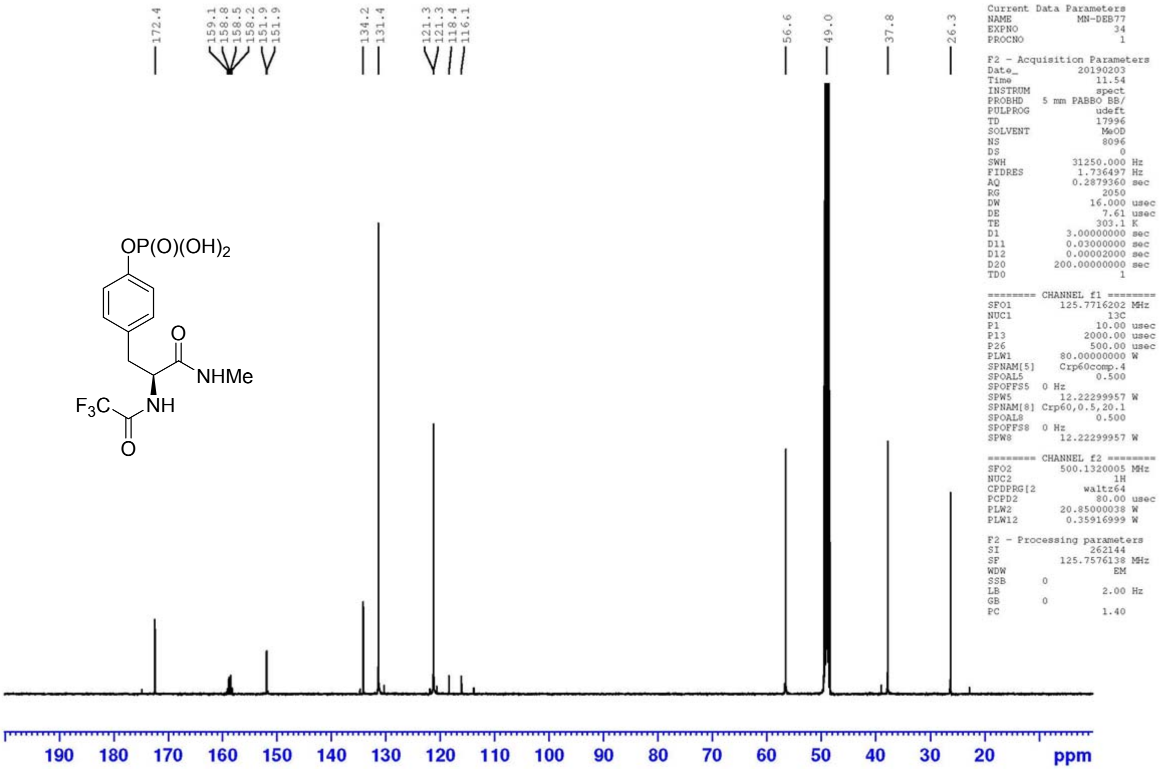

(c)

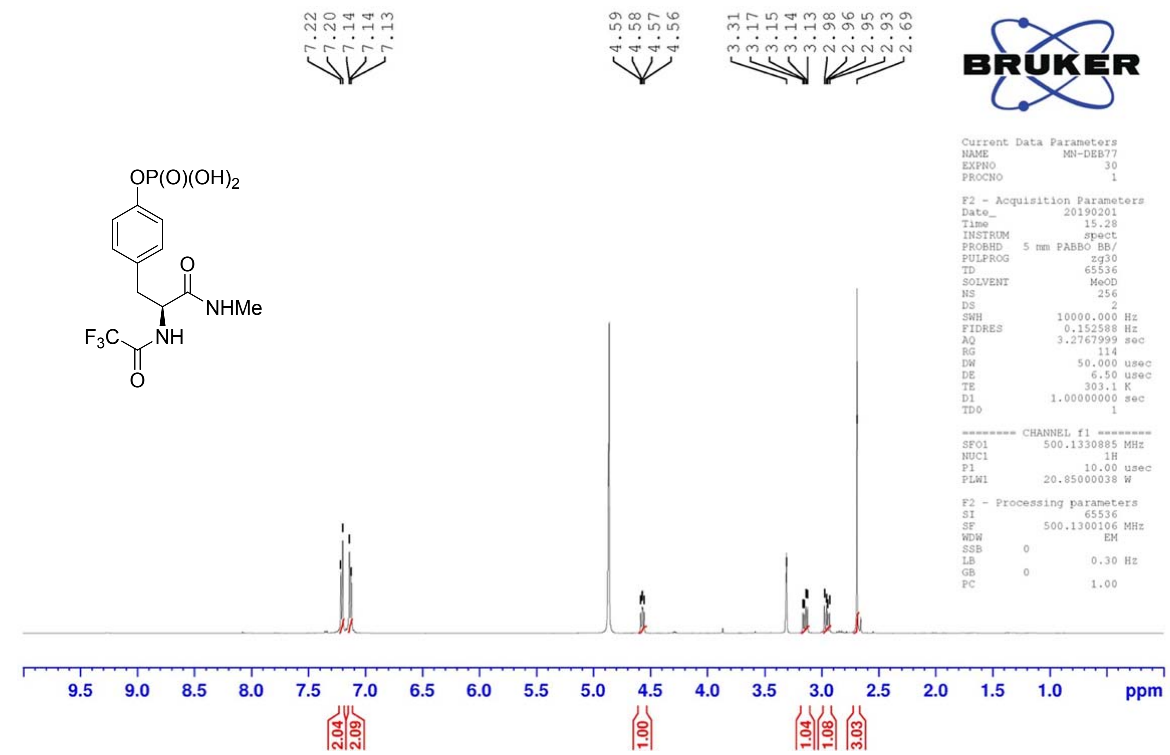


(d)

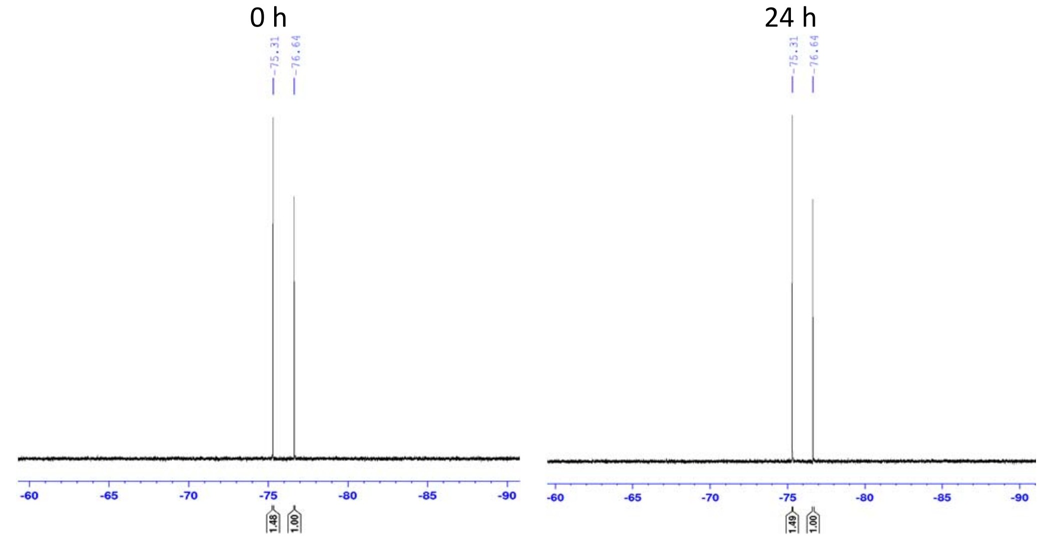


**Figure S4. Characterization of phosphate 3**

(a) Binding affinity of the phosphate 3 to SBC determined by ITC. The ITC measurement was carried out at 298K. Values reported are the means ± s.e.m. from four independent experiments (b) $^{13}$C NMR (500 MHz, $CD_3OD$) of phosphate **3** (c) $^{1}$H NMR (500 MHz, $CD_3OD$) of phosphate **3** (d) Phosphate **3** stability test. The two peaks are phosphate **3** and an internal reference trifluoroethanol in NMR buffer (20mM HEPES, pH8, 50mM NaCl and 1mM DTT) measured at 0 and 24 h at room temperature.

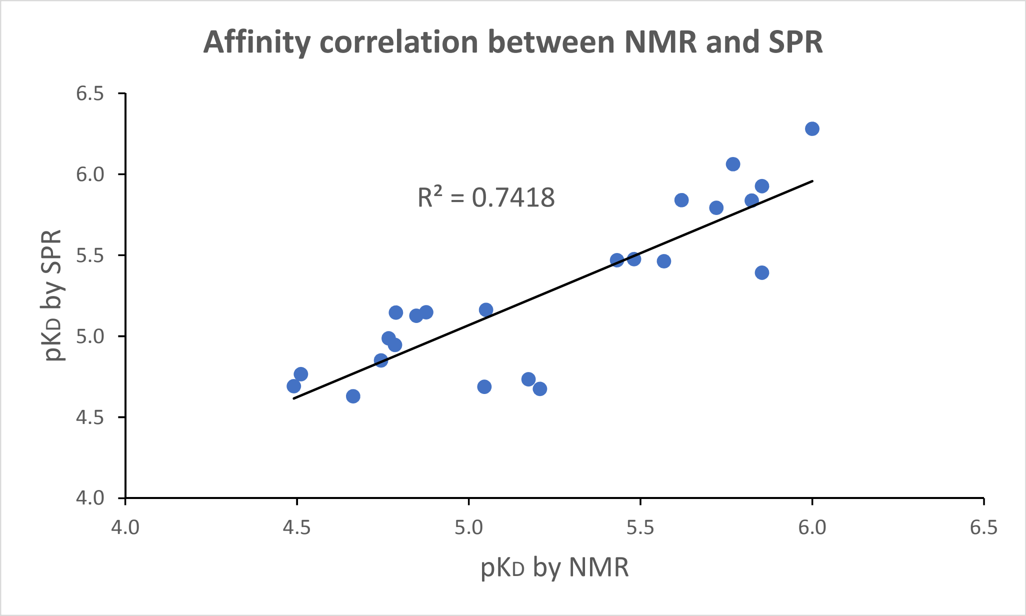


**Figure S5. Correlation of binding affinity of the alanine peptide library measured by $^{19}$F-NMR and SPR technique**

Spectra from 10 to –3 ppm

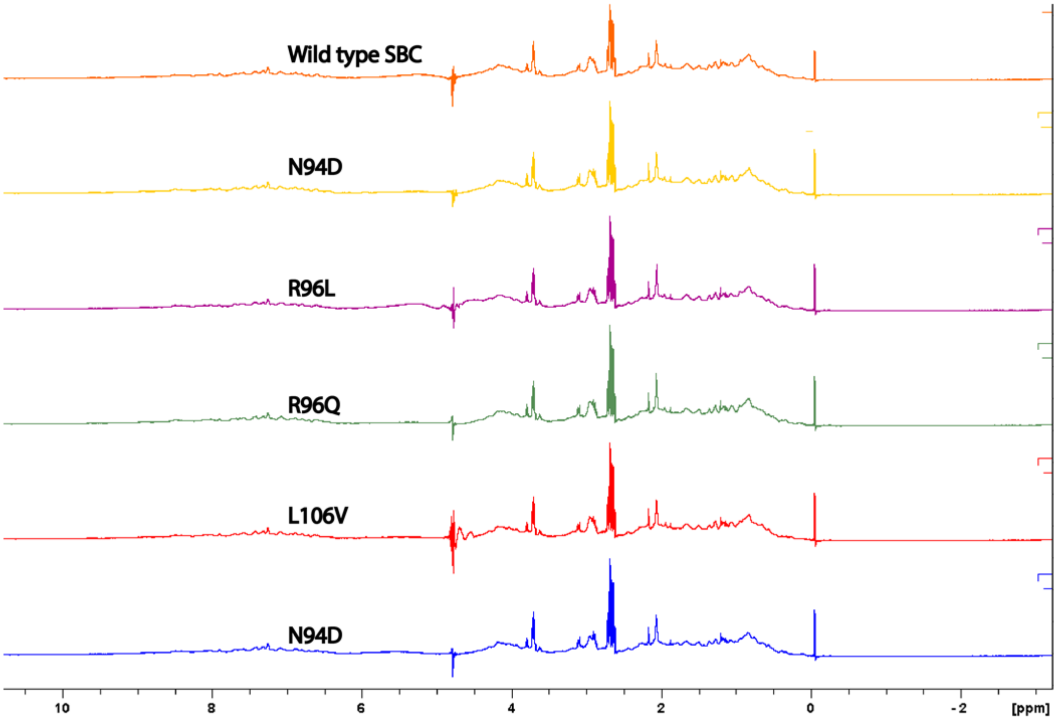

Spectra zoom in from 10 to 6 ppm

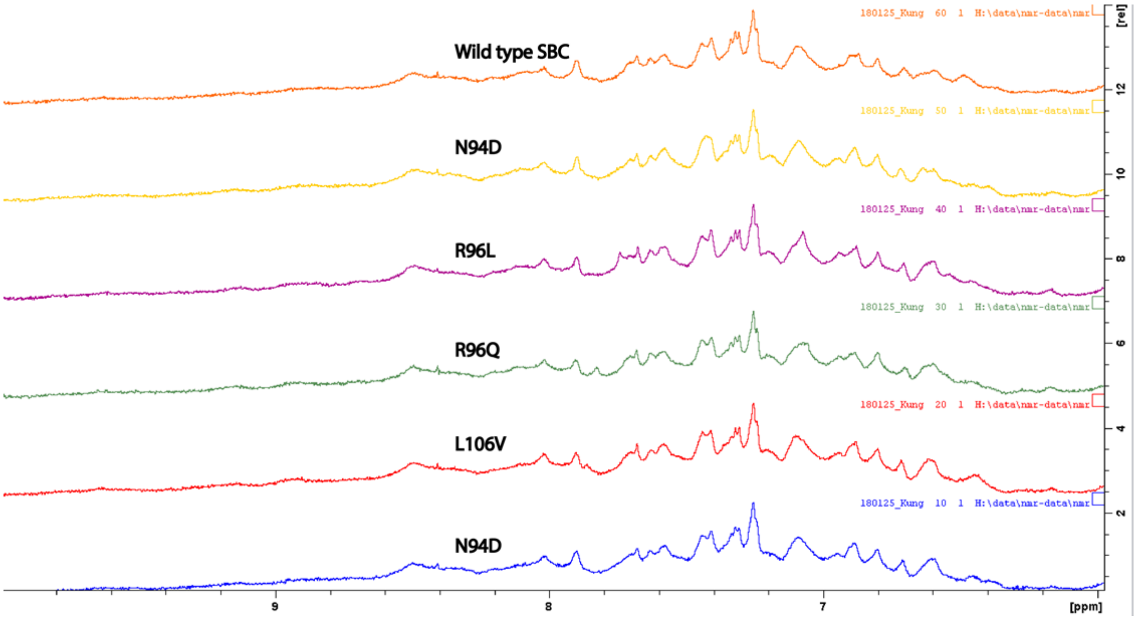


Spectra zoom in from 1.5 to –0.5 ppm

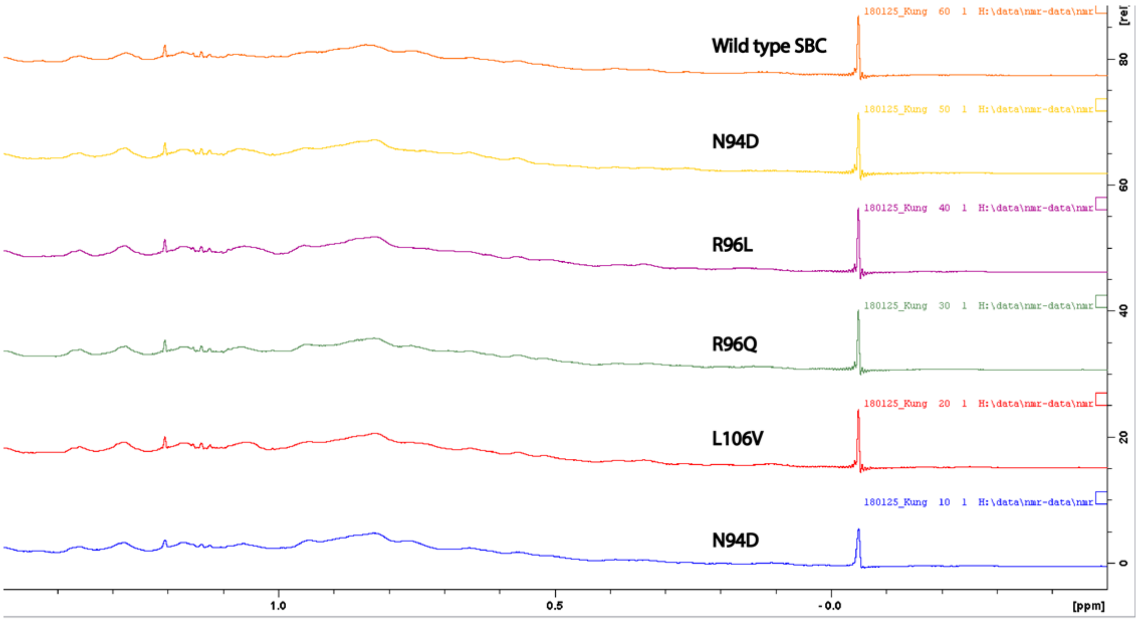


**Figure S6. The protein folding of SNPs containing SOCS2 confirmed by $^{1}$H 1D NMR**

**AVPDpYTSIHIV (MW~1293.581)**

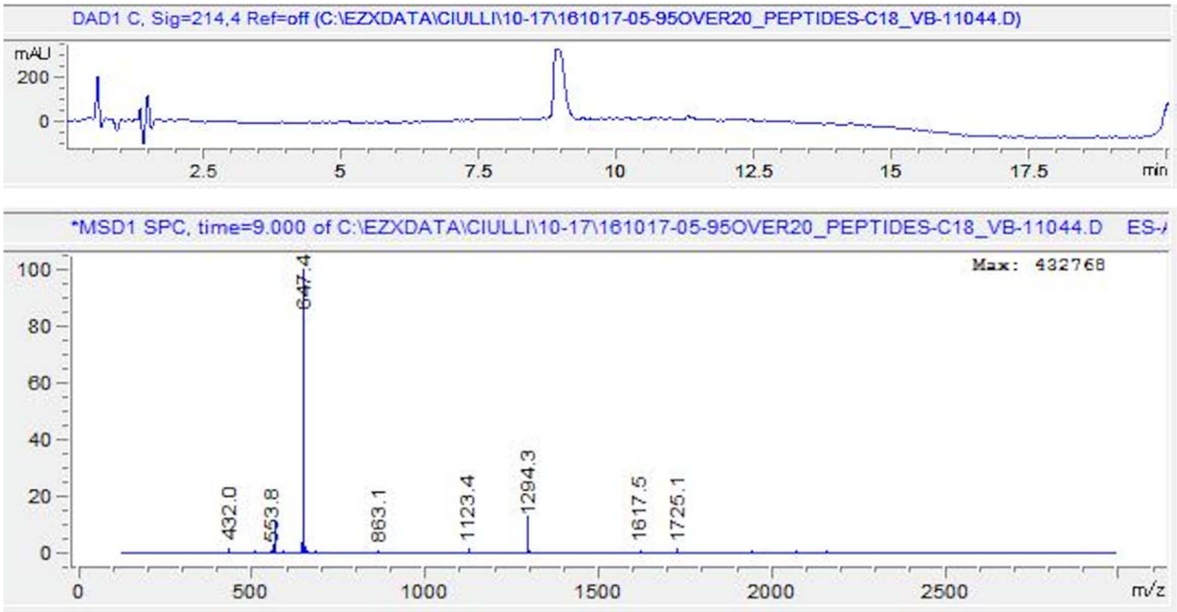


**PAPDpYTSIHIV (MW~1291.565)**

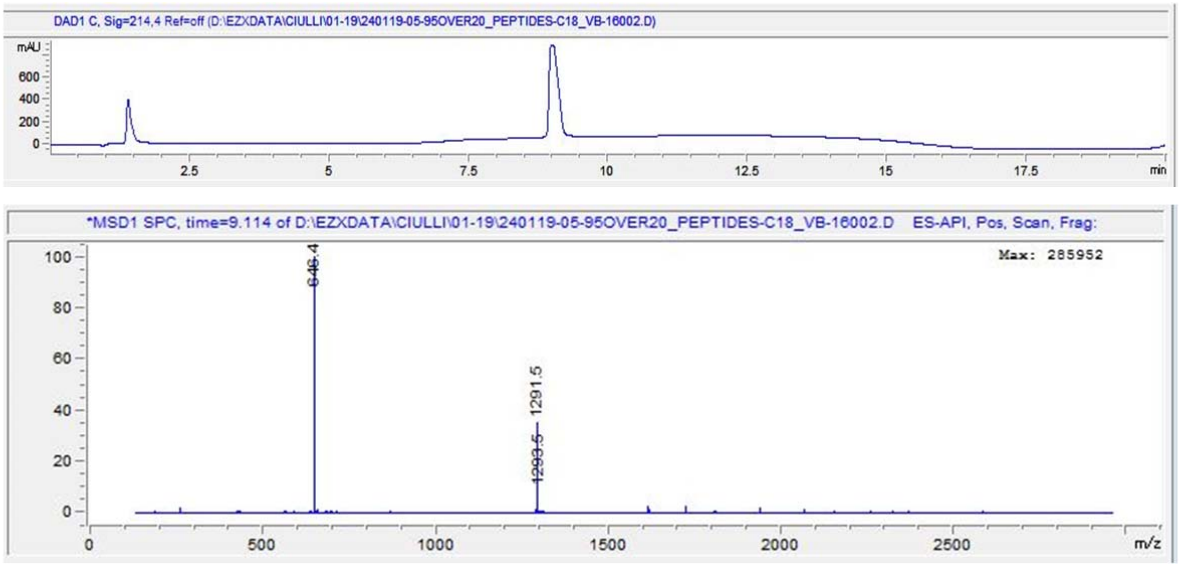

**PVADpYTSIHIV (MW~1293.581)**

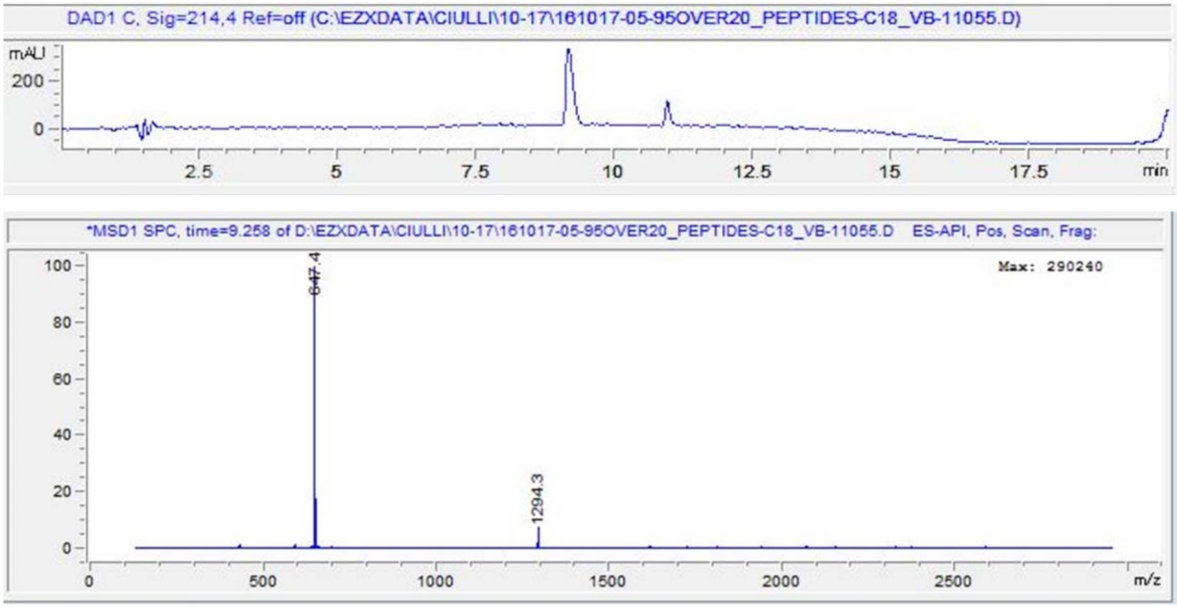


**PVPApYTSIHIV (MW~1275.607)**

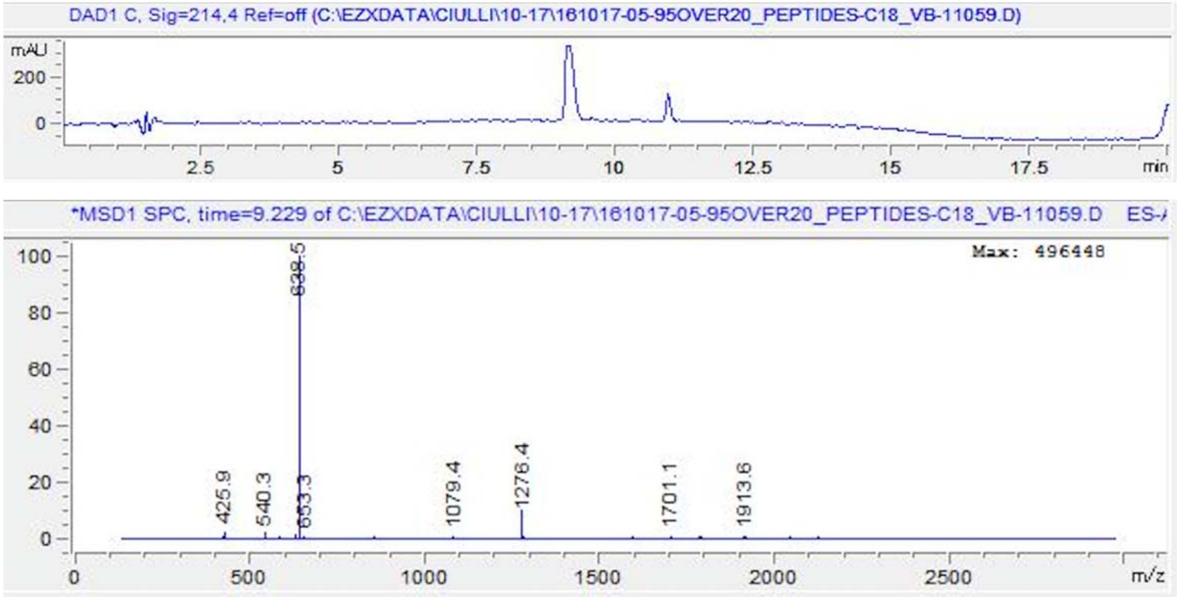

**PVPDpYASIHIV (MW~1289.586)**

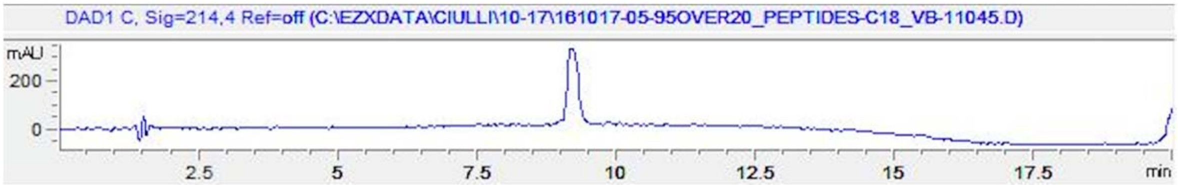


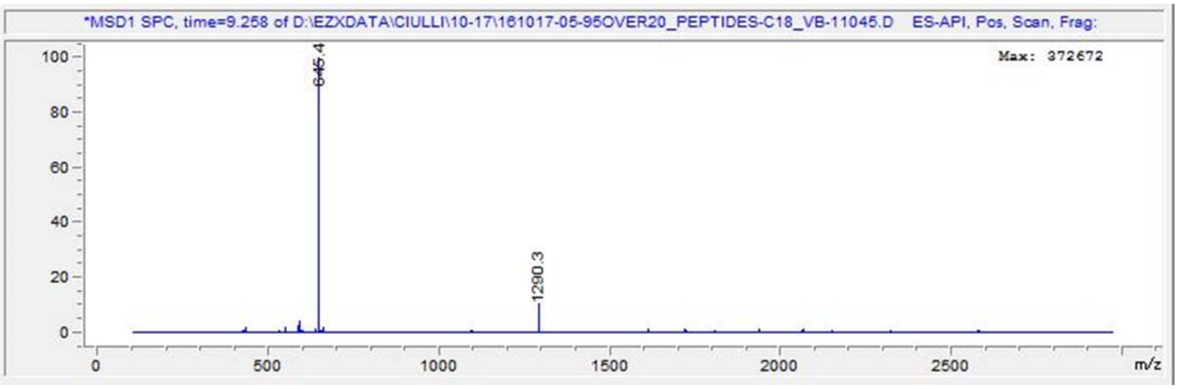


**PVPDpYTAIHIV (MW~1303.602)**

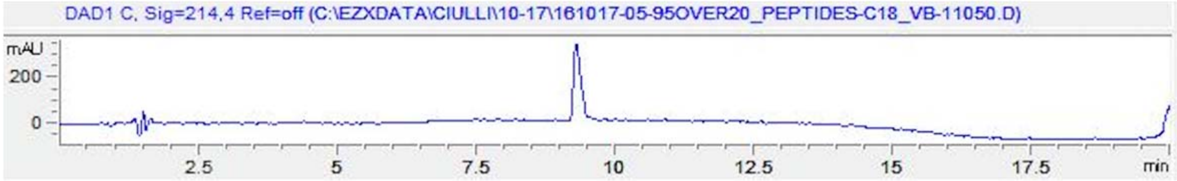


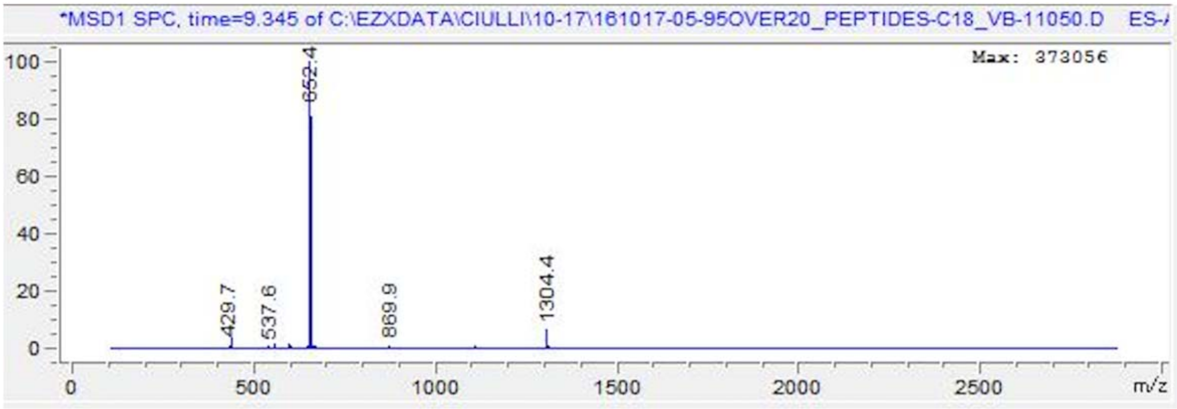

**PVPDpYTSAHIV (MW~1277.550)**

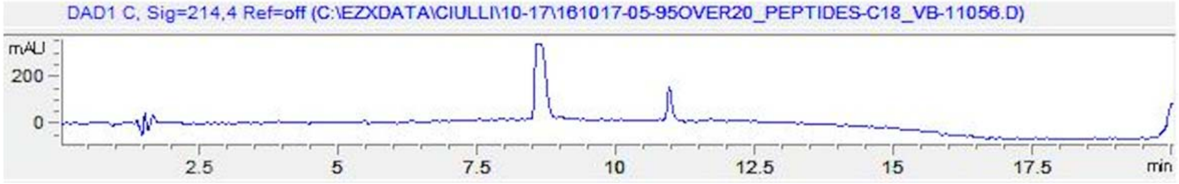


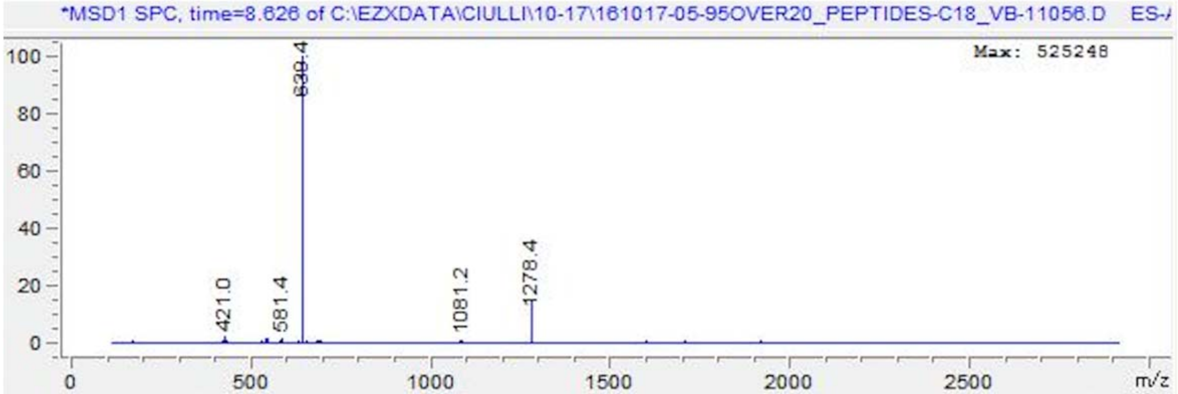


**PVPDpYTSIAIV (MW~1253.575)**

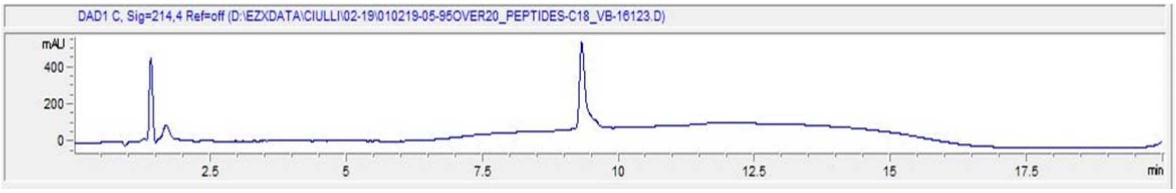


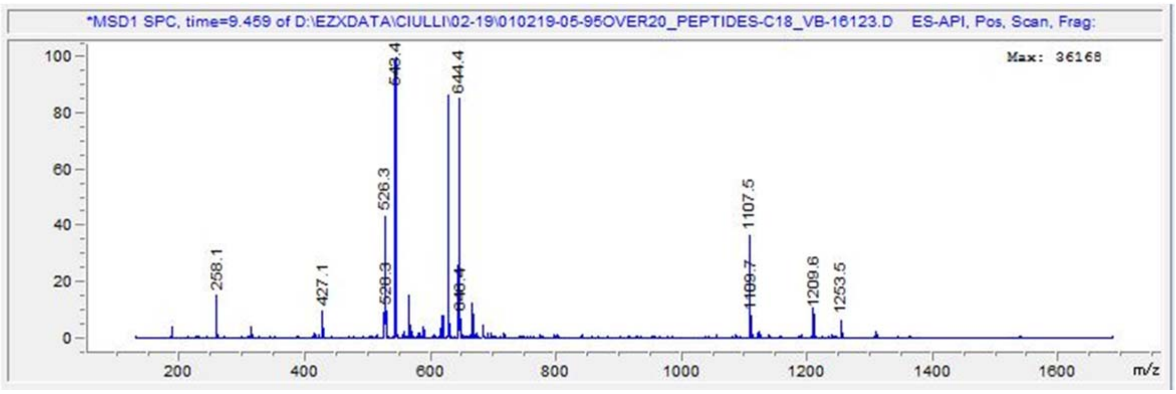

**PVPDpYTSIHAV (MW~1277.550)**

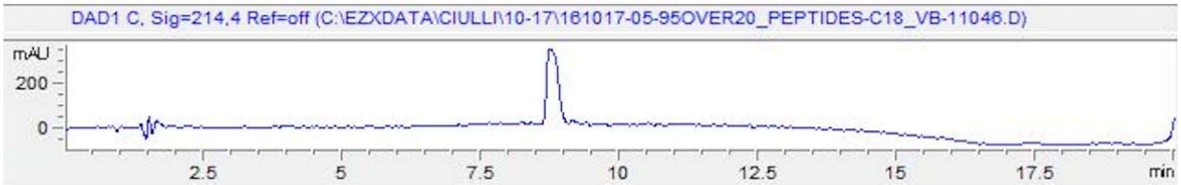


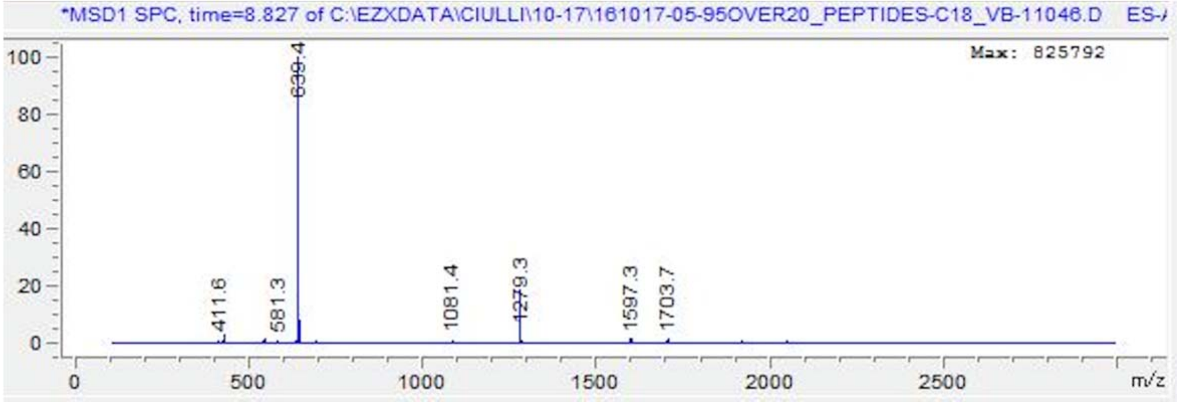


**PVPDpYTSIHIA (MW~1291.565)**

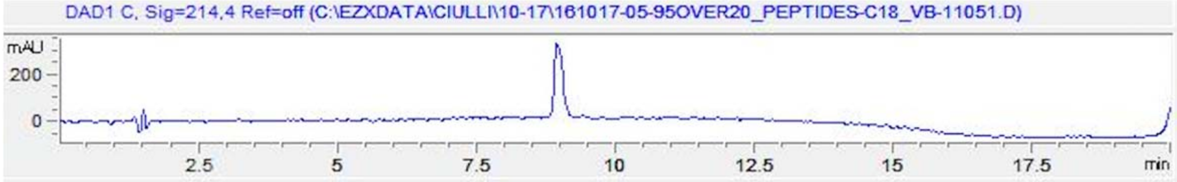


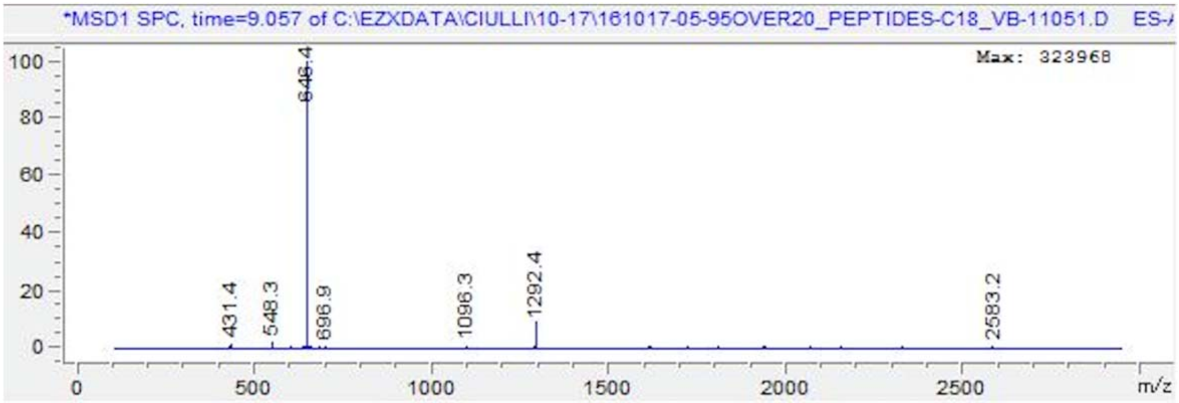

**ASFEpYTILDPS (MW~1321.528)**

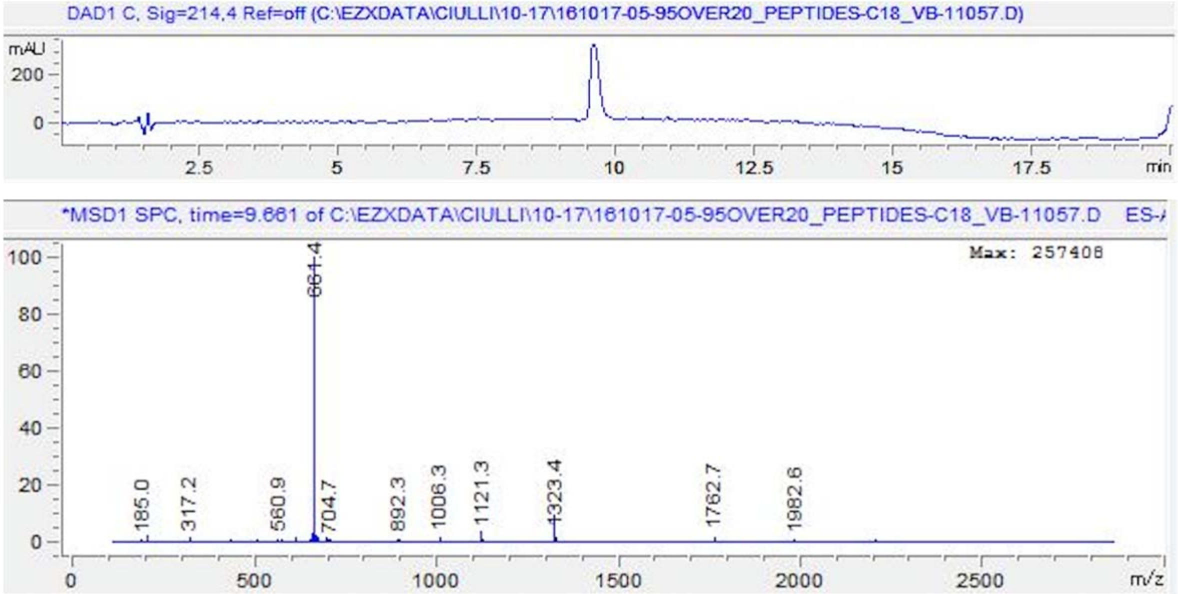


**AAFEpYTILDPS (MW~1305.533)**

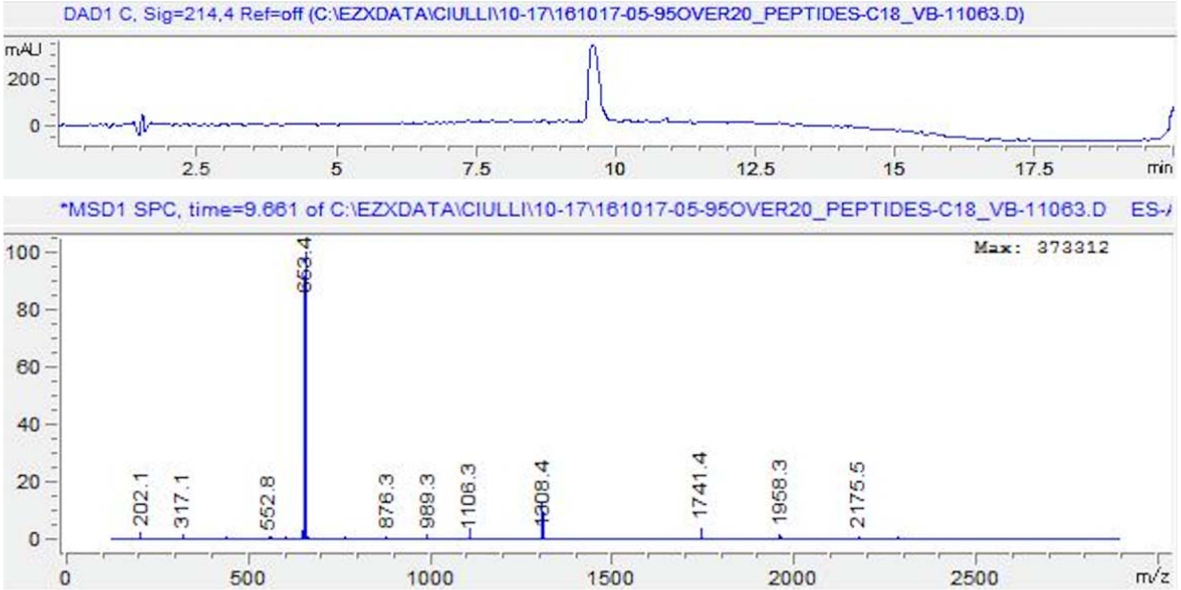

**ASAEpYTILDPS (MW~1245.497)**

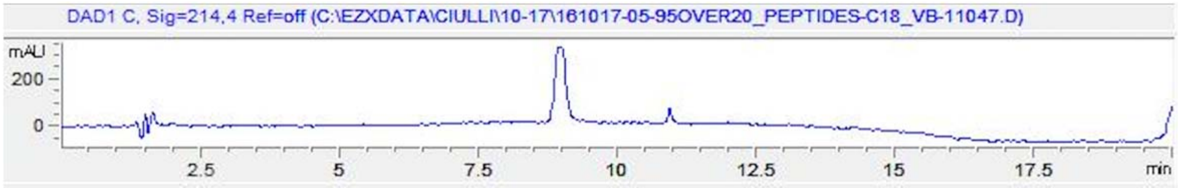


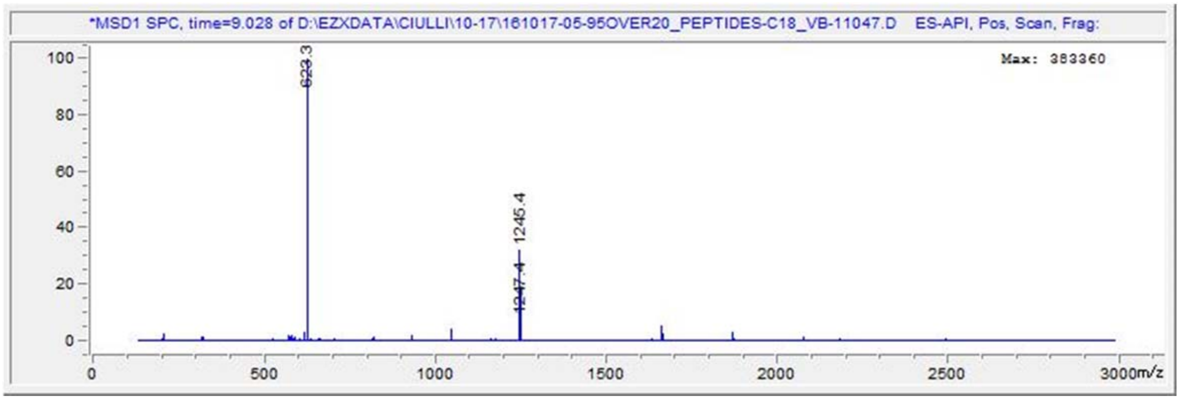


**ASFApYTILDPS (MW~1263.523)**

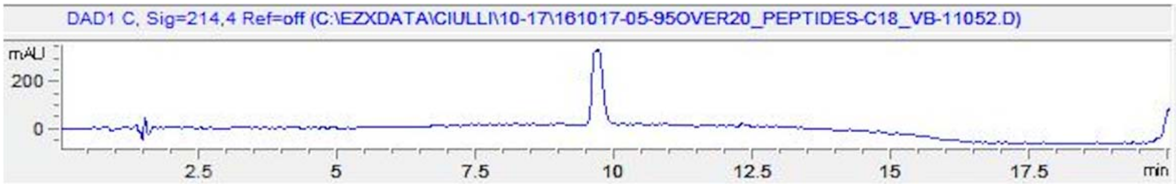


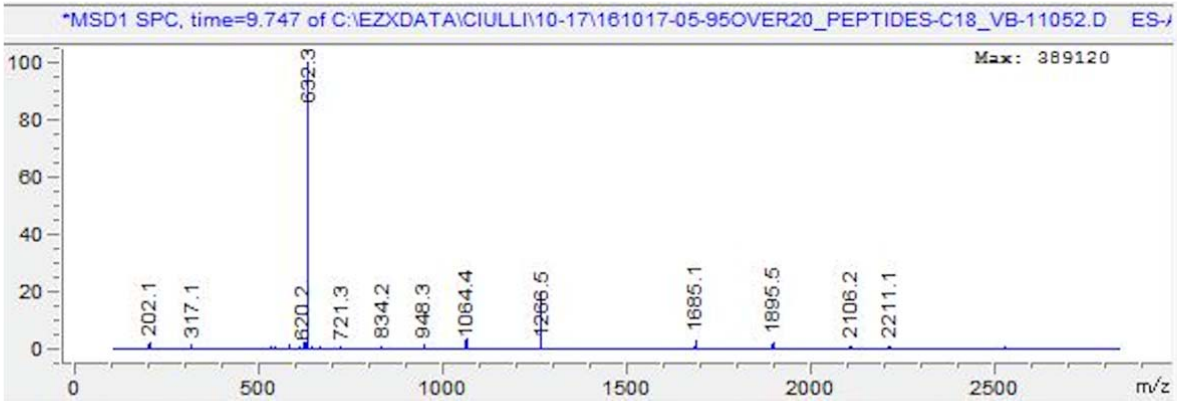

**ASFEpYAILDPS (MW~1291.518)**

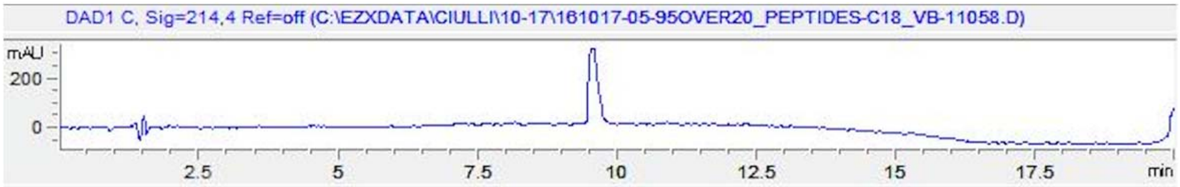


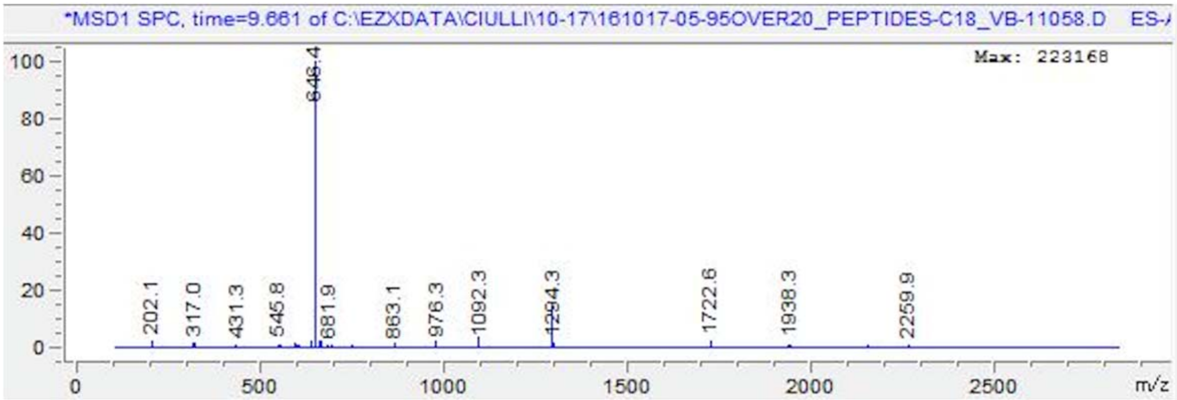


**ASFEpYTALDPS (MW~1279.481)**

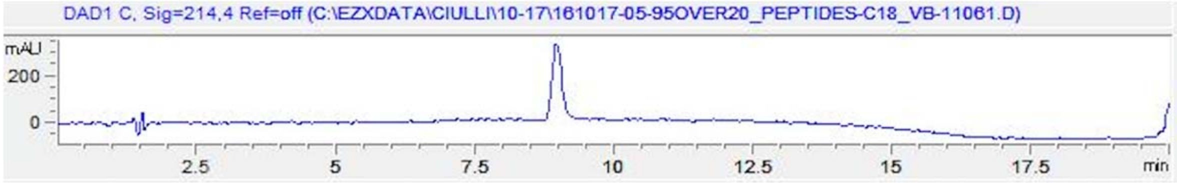


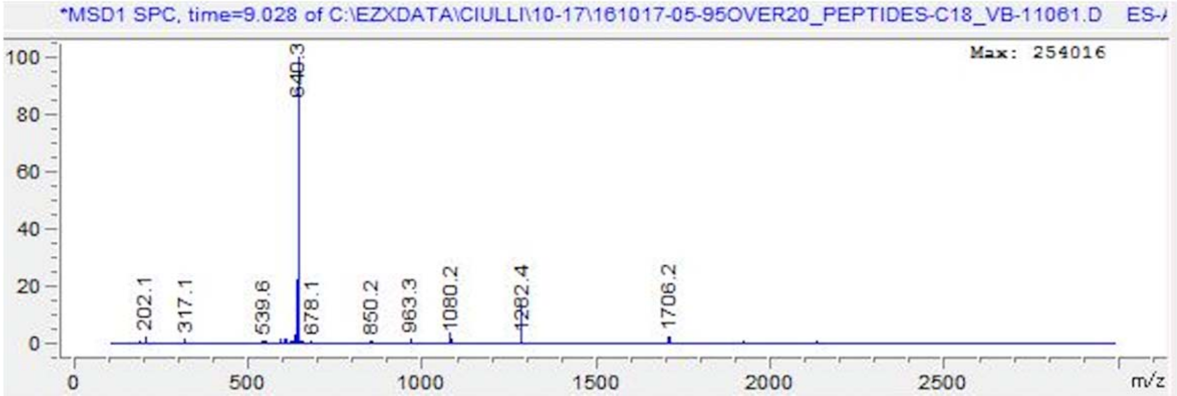

**ASFEpYTIADPS (MW~1279.481)**

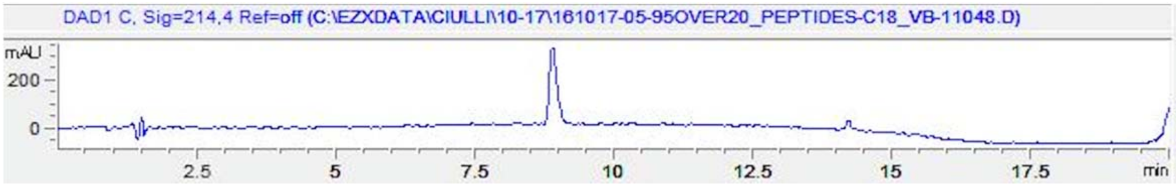


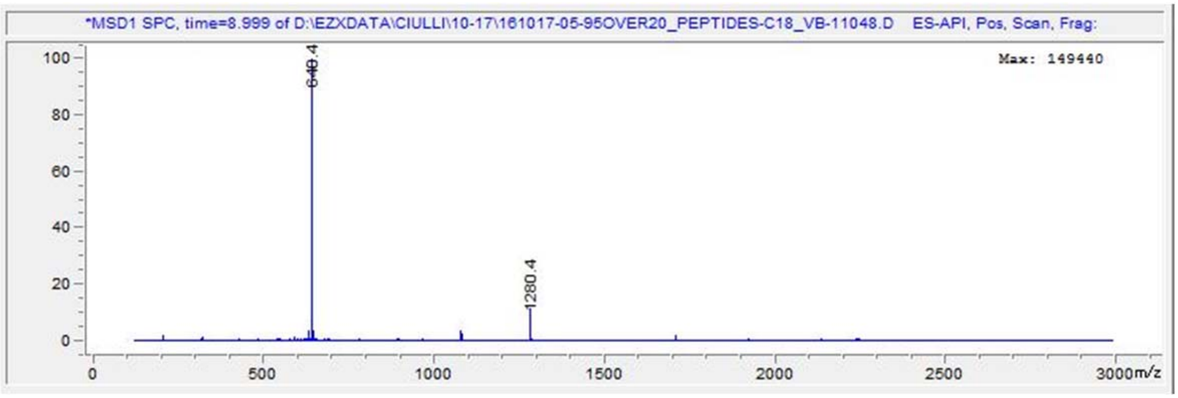


**ASFEpYTILAPS (MW~1277.538)**

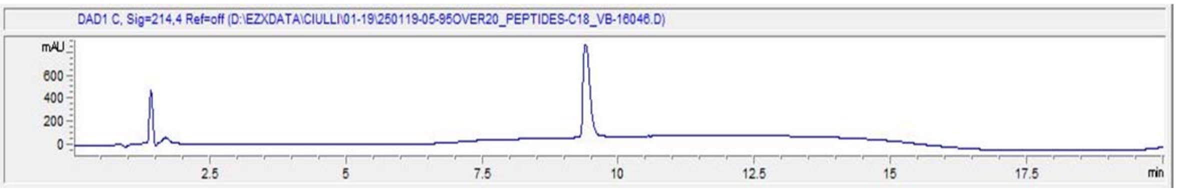


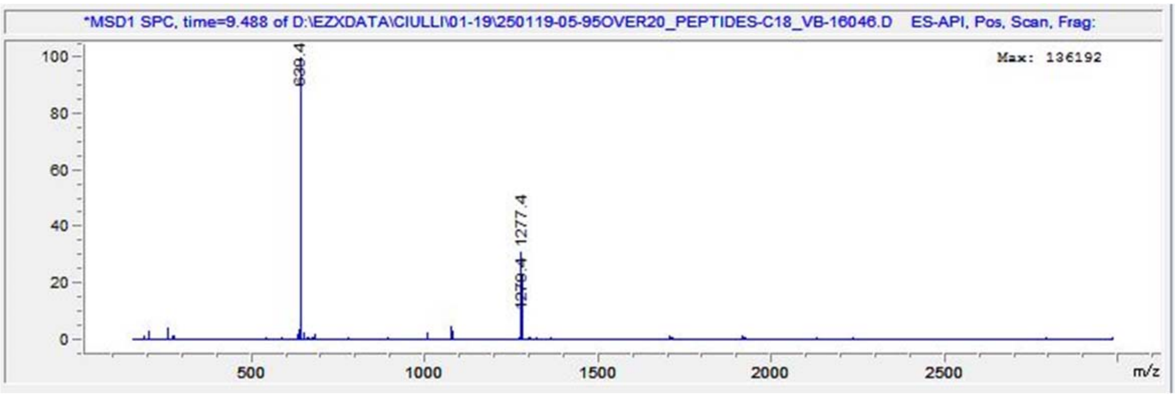

**ASFEpYTILDAS (MW~1295.513)**

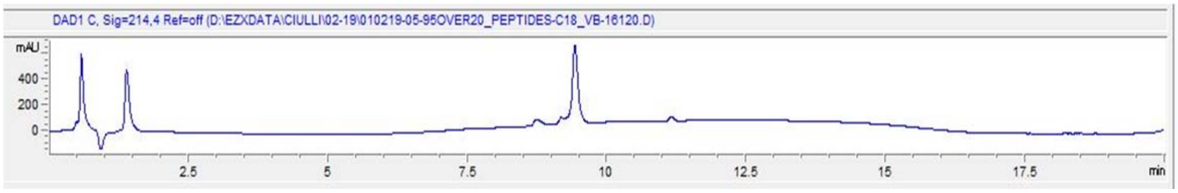


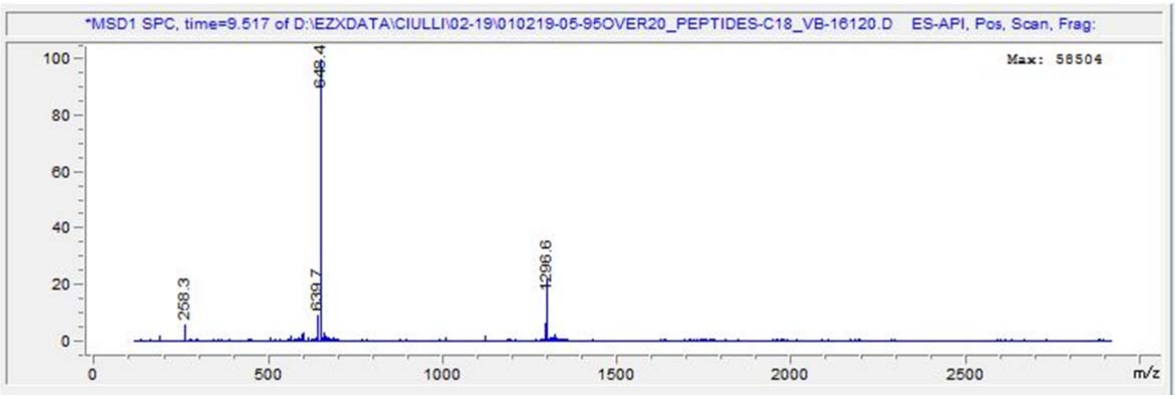


**ASFEpYTILDPA (MW~1305.533)**

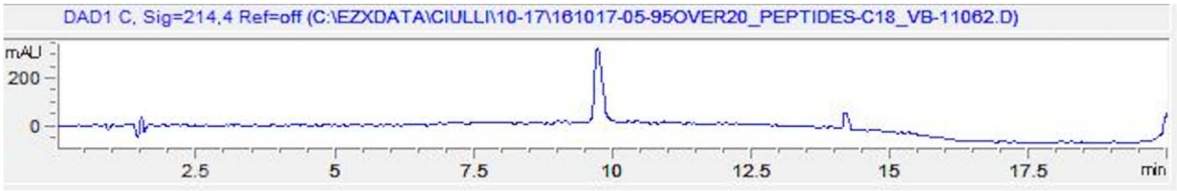


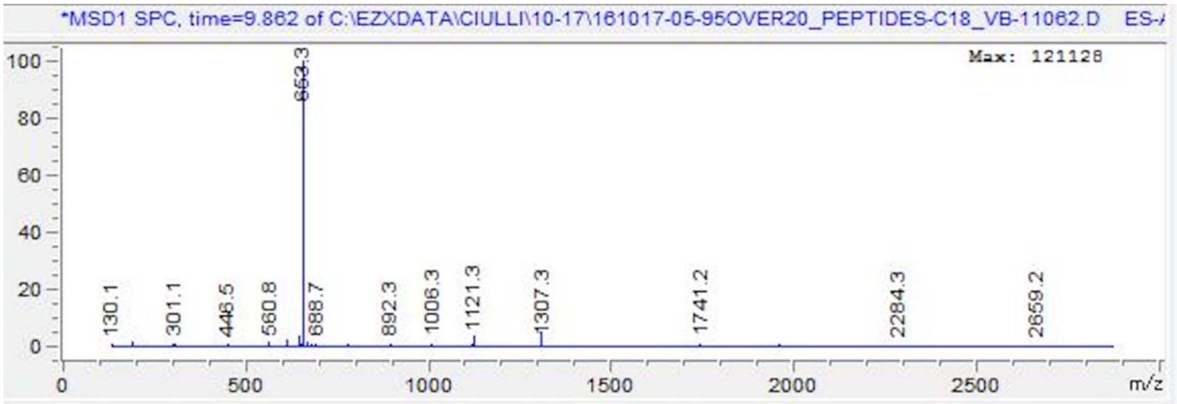

**GHR mutant V(-3)R**

**PRPDpYTSIHIV (MW~1376.629)**

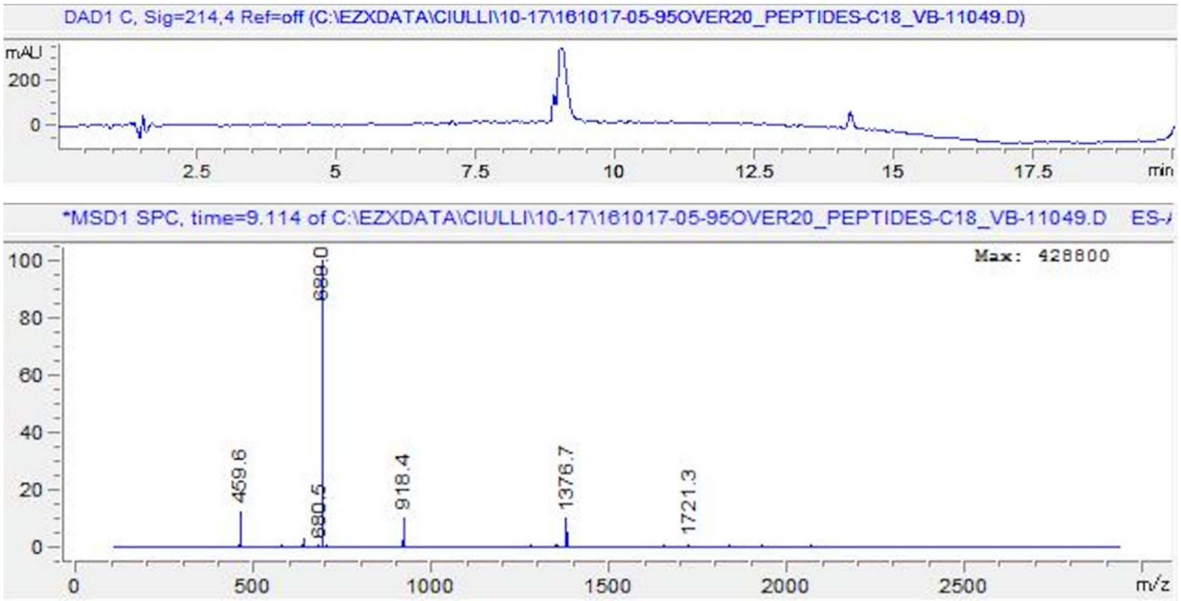


**GHR mutant V(-3)Y**

**PYPDpYTSIHIV (MW~1383.591)**

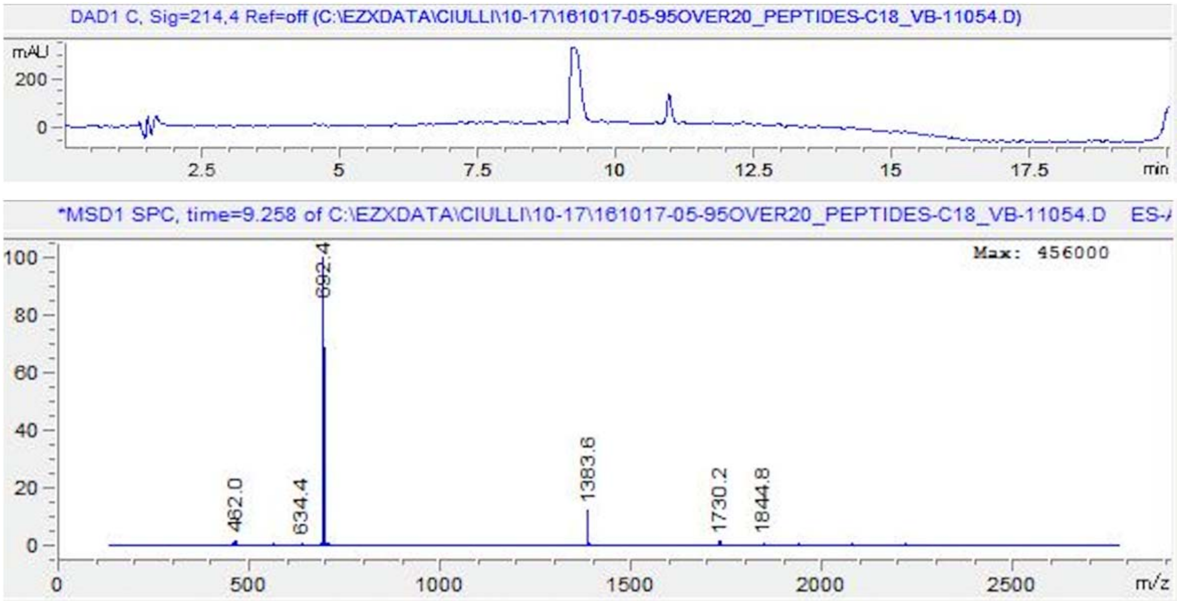

**GHR_pY487 NIDFpYAQVSDI (MW~1363.550)**

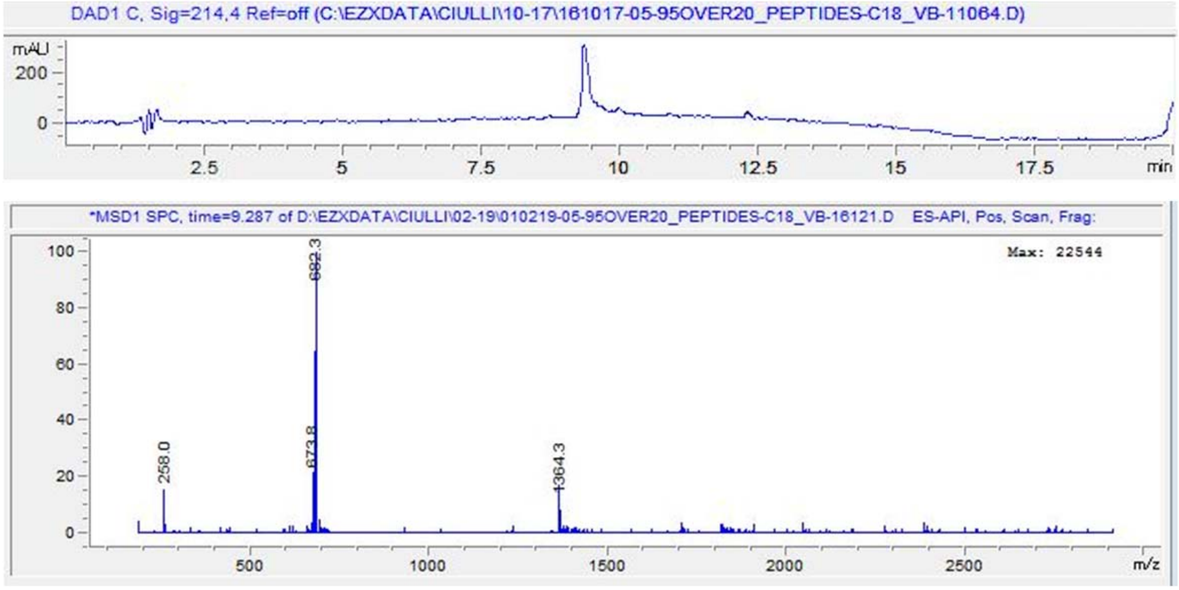


**GHR_pY595 PVPDpYTSIHIV (MW~1319.597)**

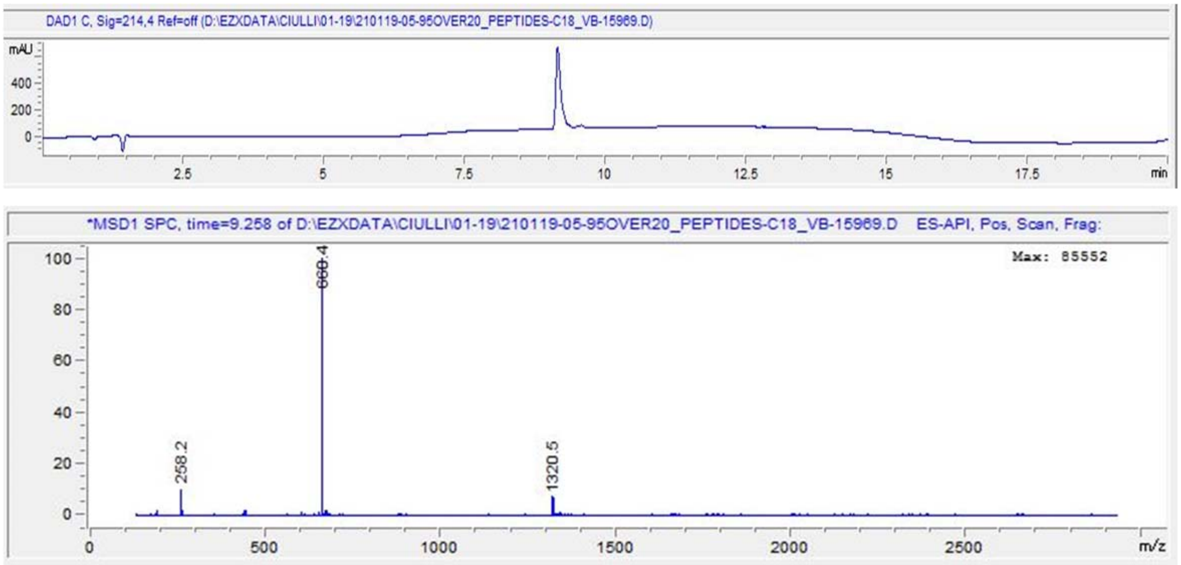


**Figure S7. The LC-MS data of synthesized peptide**

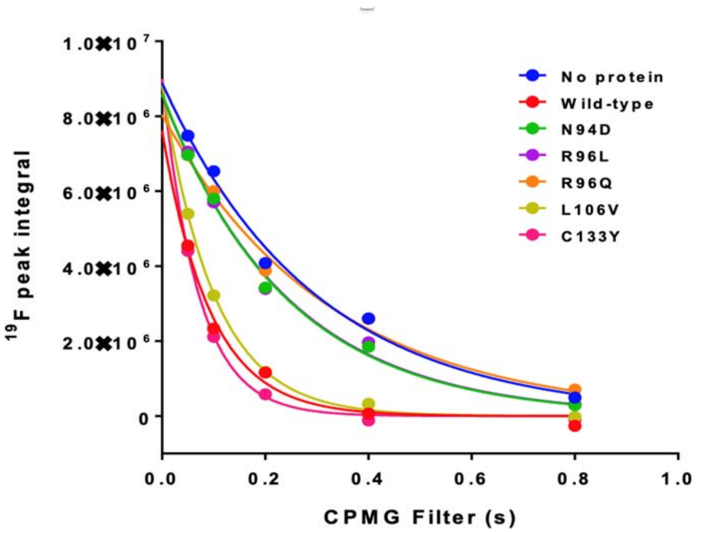


| One phase decay | No protein | Wild-type | N94D | R96L | R96Q | L106V | C133Y |
|---|---|---|---|---|---|---|---|
| Best-fit values | | | | | | | |
| Y0 | 8885000 | 7611000 | 8593000 | 8572000 | 8040000 | 8857000 | 8997000 |
| K | 3.402 | 10.75 | 4.179 | 4.136 | 3.114 | 9.999 | 14.33 |
| Tau | 0.294 | 0.09303 | 0.2393 | 0.2418 | 0.3212 | 0.1 | 0.06981 |
| Std. Error | | | | | | | |
| Y0 | 418918 | 920328 | 336002 | 404716 | 369681 | 305191 | 545388 |
| K | 0.3361 | 1.631 | 0.3177 | 0.3811 | 0.3094 | 0.45 | 0.92 |
| R square | 0.9899 | 0.9853 | 0.9946 | 0.9919 | 0.9889 | 0.9986 | 0.9975 |

**Figure S8. Measurement of the transverse relaxation rate ($R_2$) of the spy in the absence and presence of proteins. Data fitted by prism.**

Peptides were measured at concentration of 0.08, 0.25, 0.7, 2.2, 6.7 20 and 60 µM.

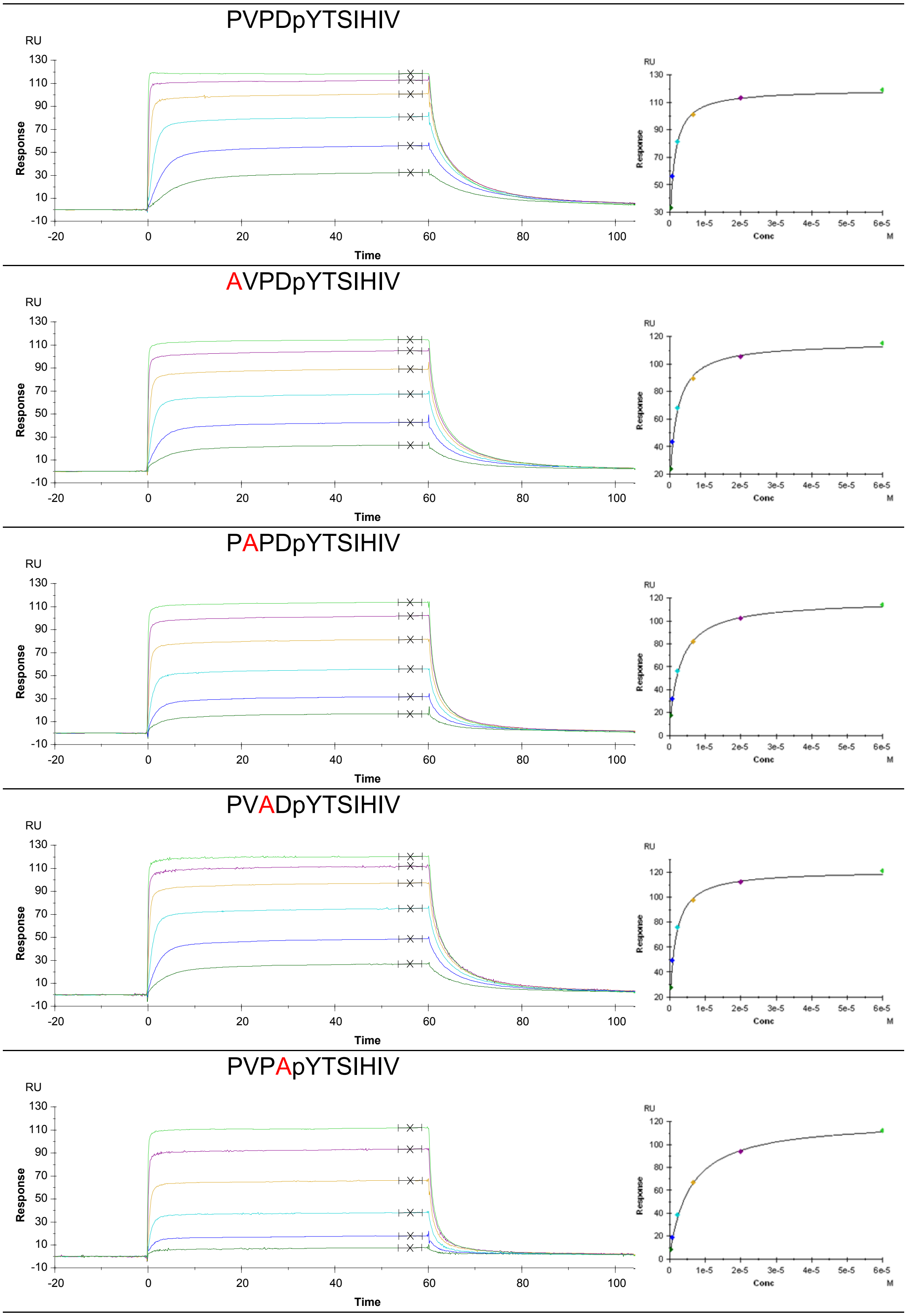

PVPDpYASIHIV

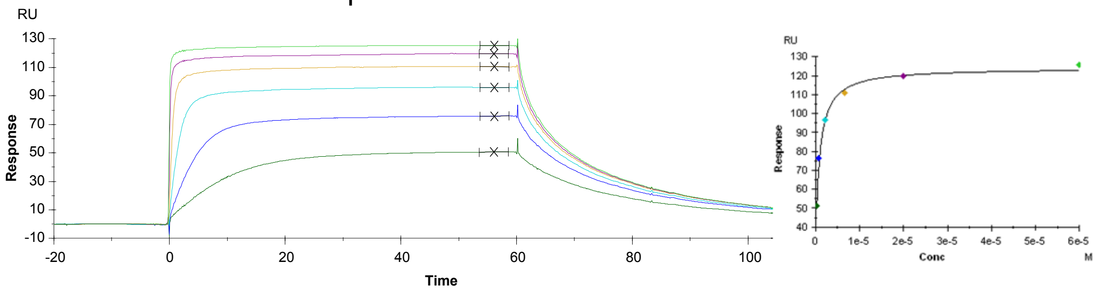


PVPDpYTAIHIV

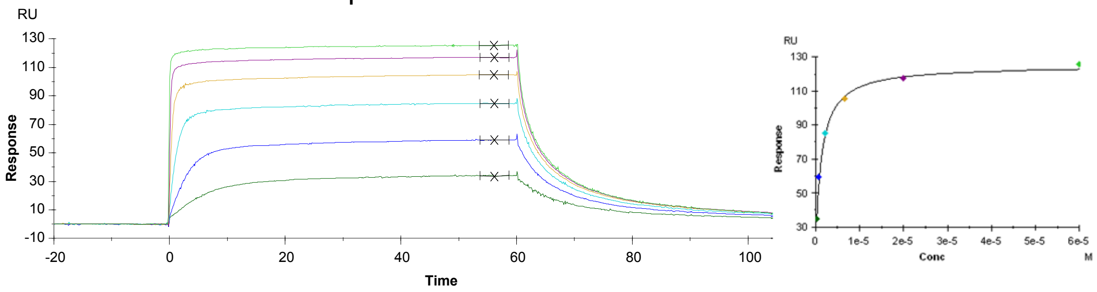


PVPDpYTSAHIV

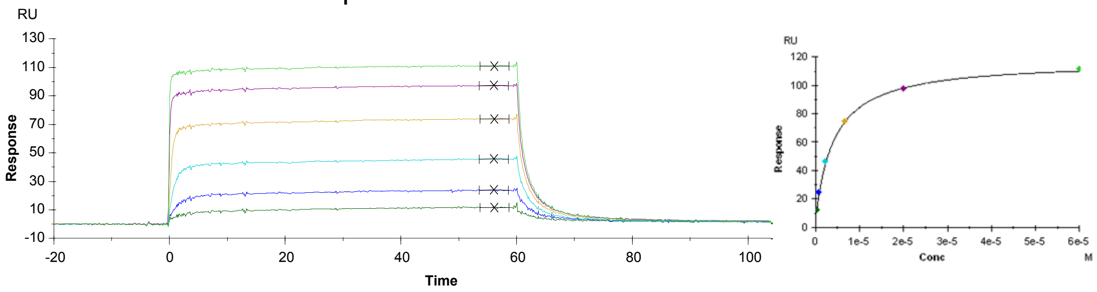


PVPDpYTSIAIV

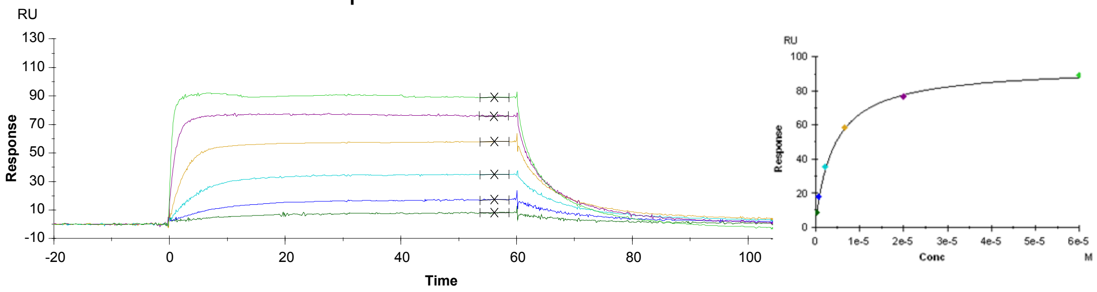


PVPDpYTSIHAV

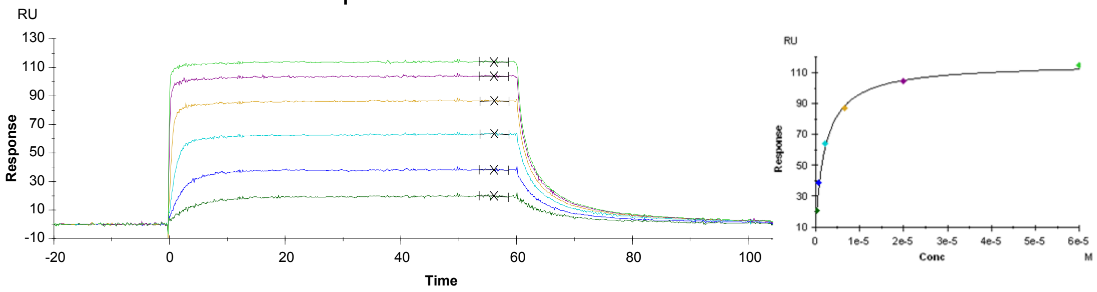

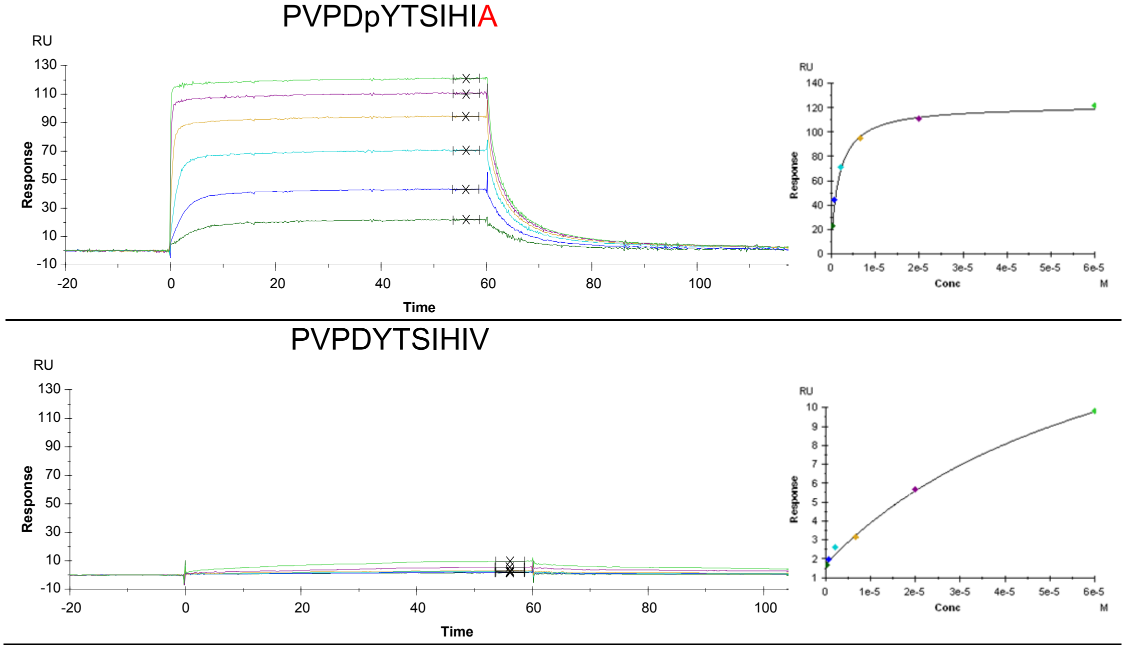


**Figure S9. Alanine scan of GHR_pY595 derivatives by SPR**

Peptides were measured at concentration of 0.08, 0.25, 0.7, 2.2, 6.7 20 and 60 μM.

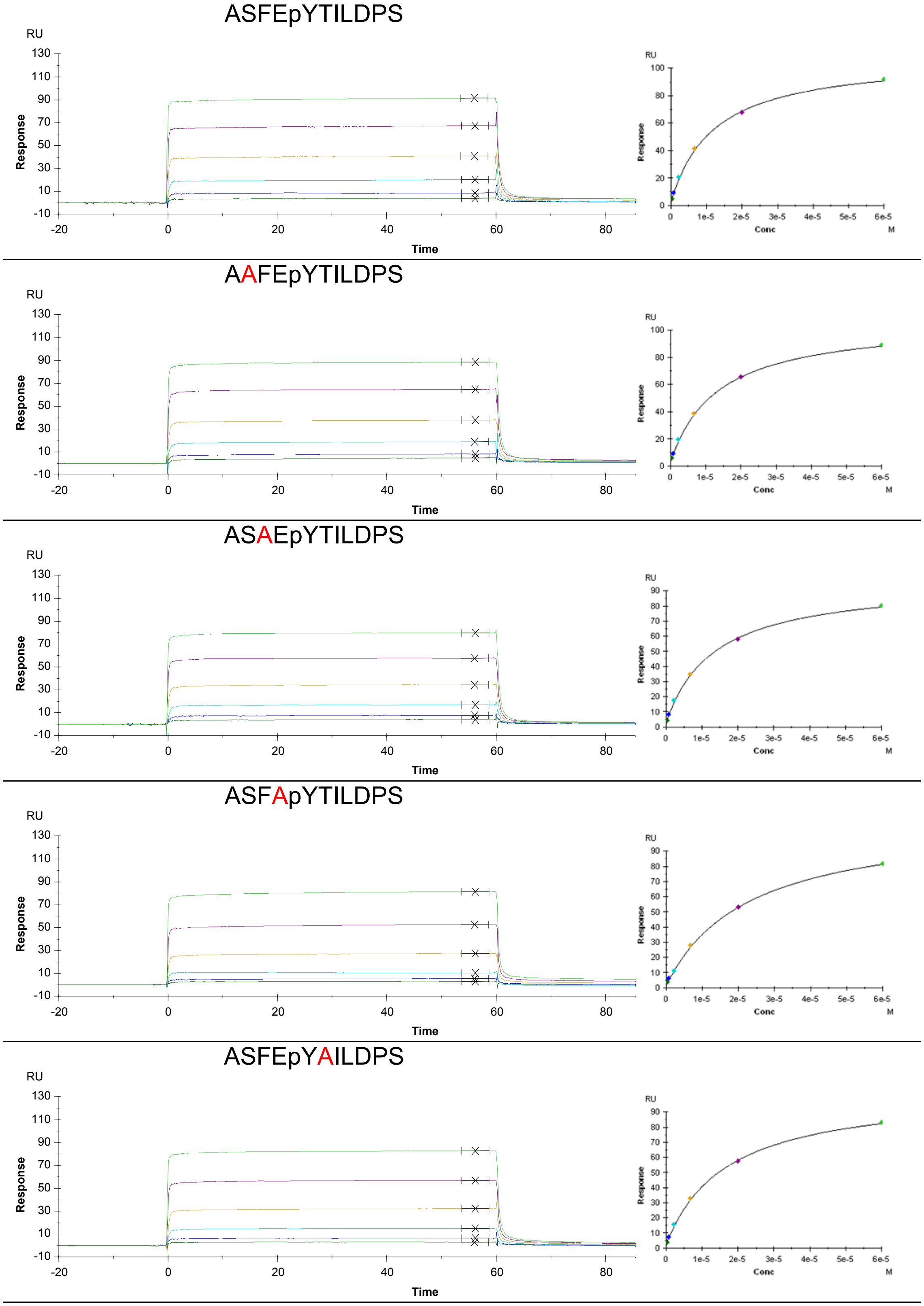

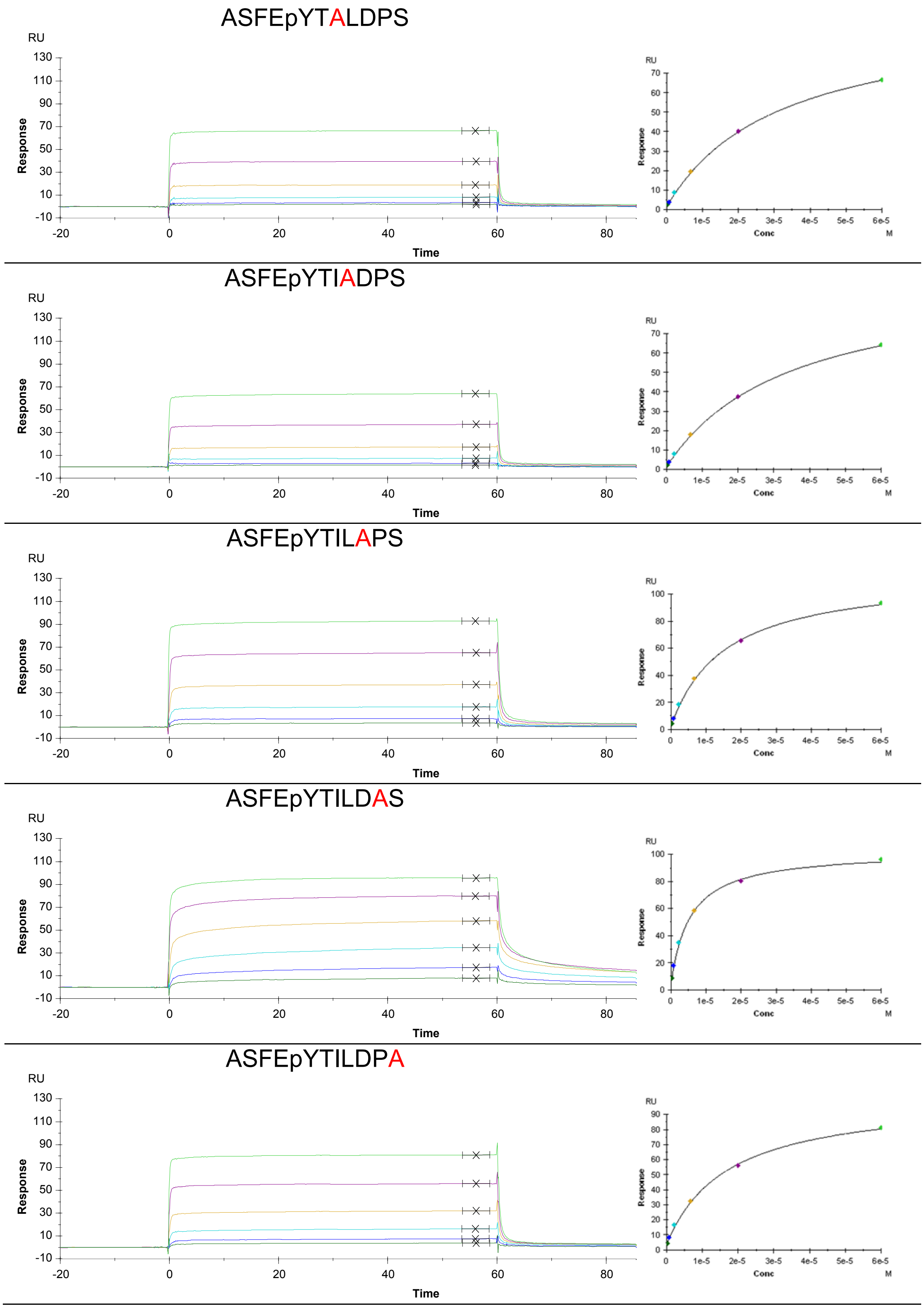


**Figure S10. Alanine scan of EpoR_pY426 derivatives by SPR**

Peptides were measured at concentration of 0.08, 0.25, 0.7, 2.2, 6.7 20 and 60 μM.

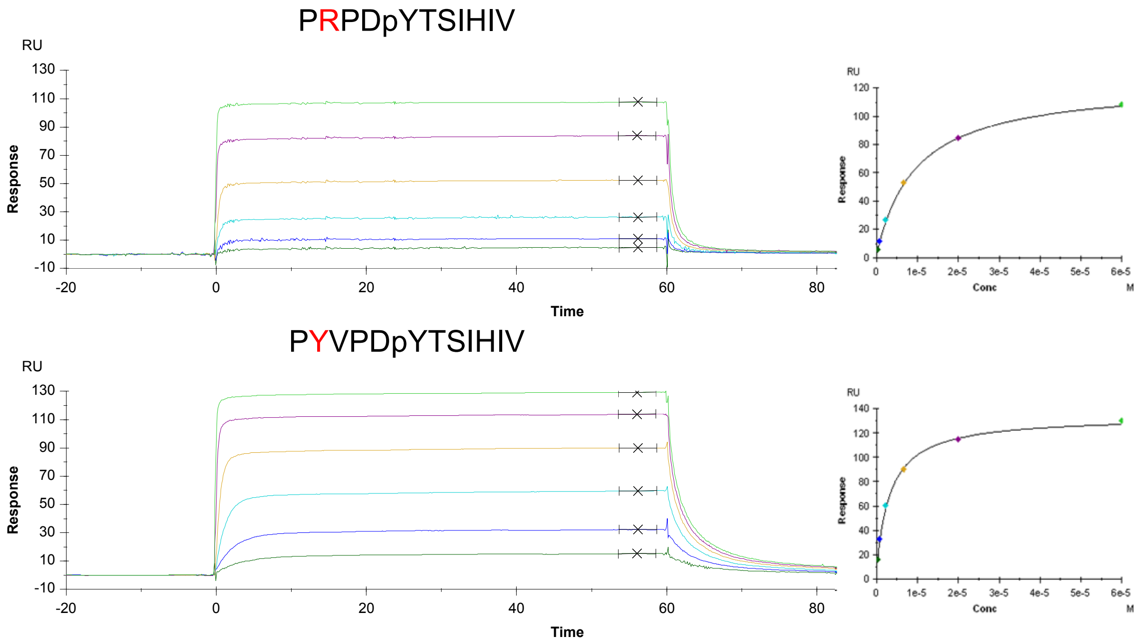


**Figure S11. $K_D$ measurement of the GHR_pY595 with Val(-3) substituted by arginine or tyrosine**

### Wild type

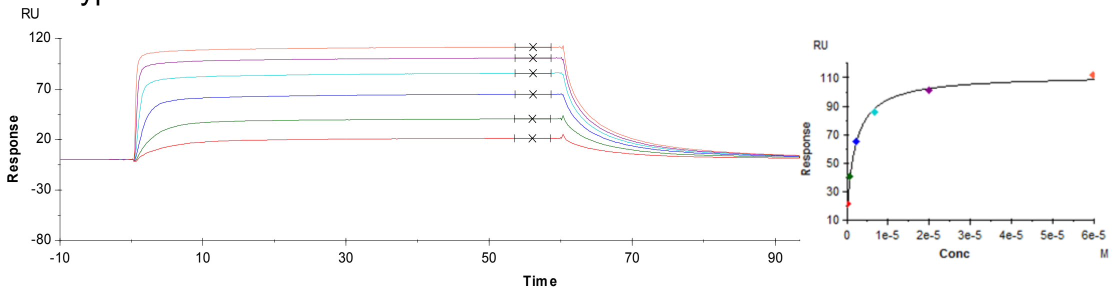


## N94D

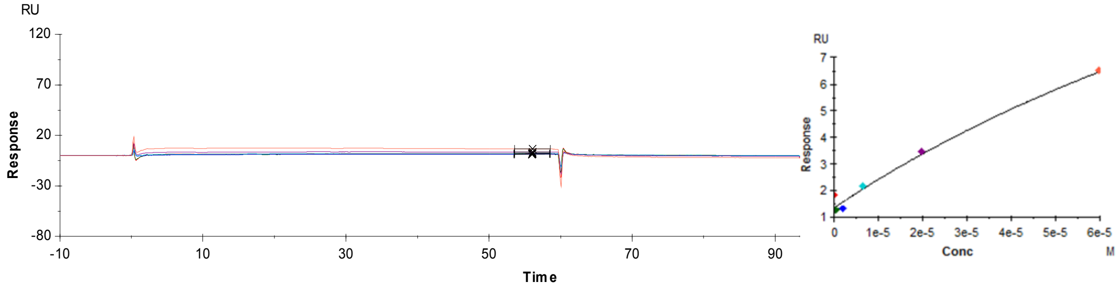


## R96L

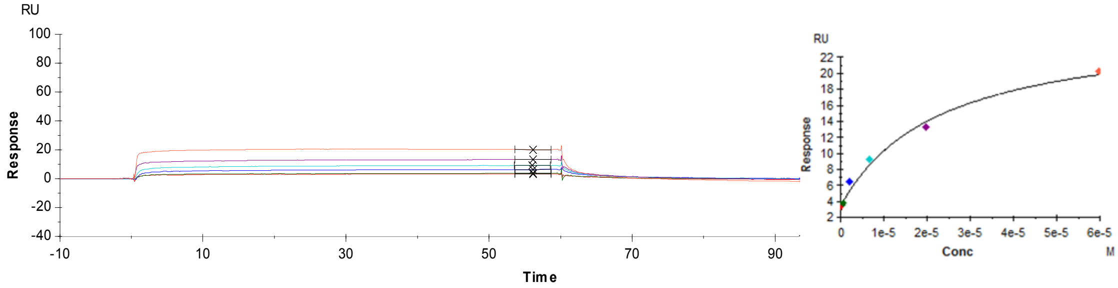


## R96Q

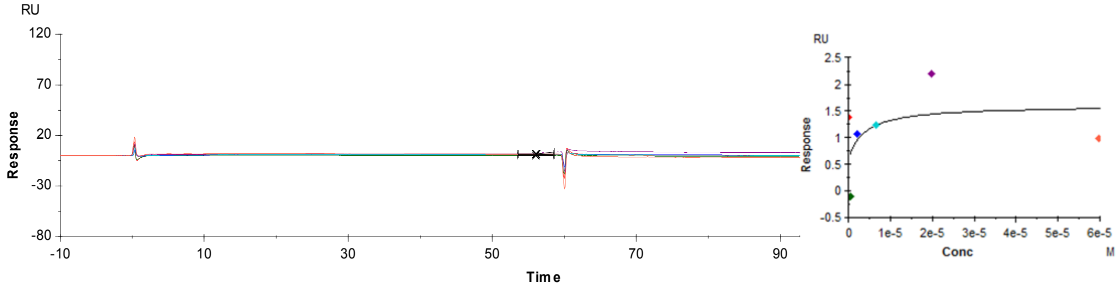


## L106V

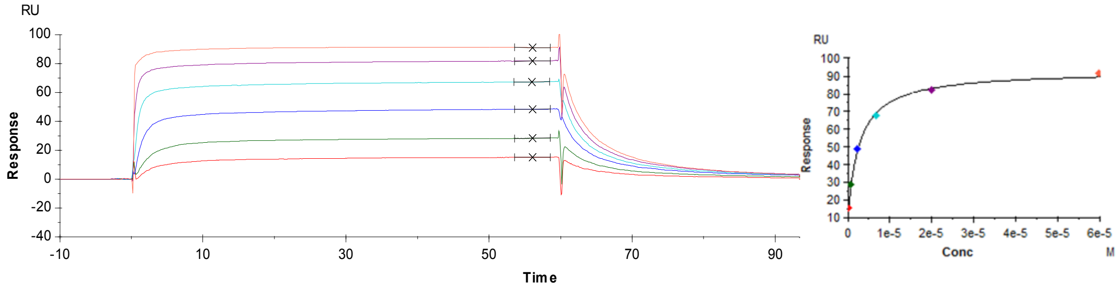


## C133Y

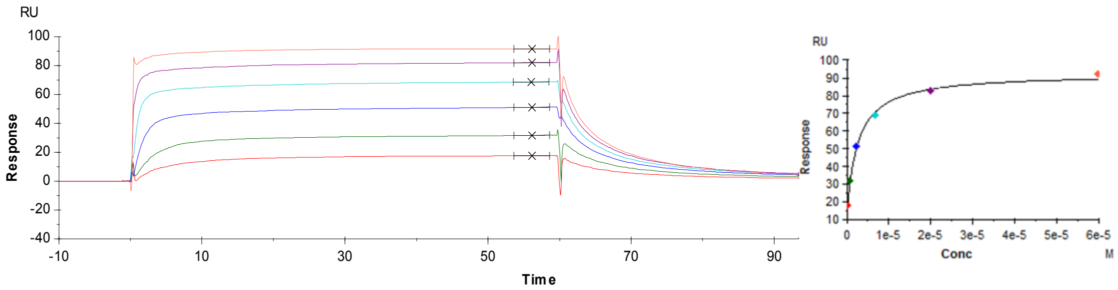


**Figure S12. The $K_D$ measurement of GHR_pY595 peptide against SNP mutants by SPR**

Peptides were measured at concentration of 0.08, 0.25, 0.7, 2.2, 6.7 20 and 60 μM.

WT

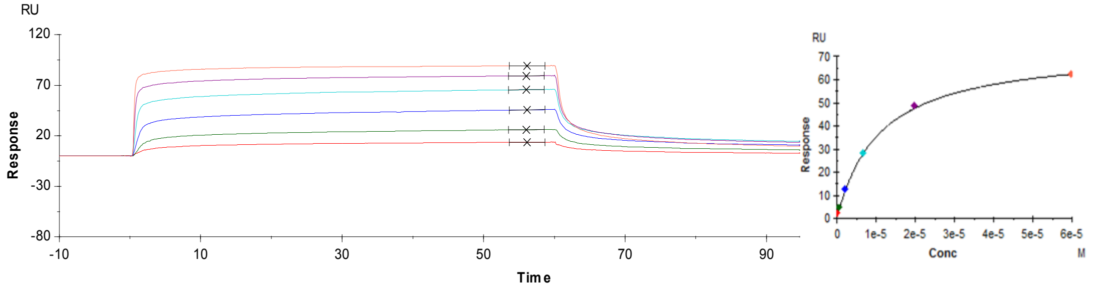


N94D

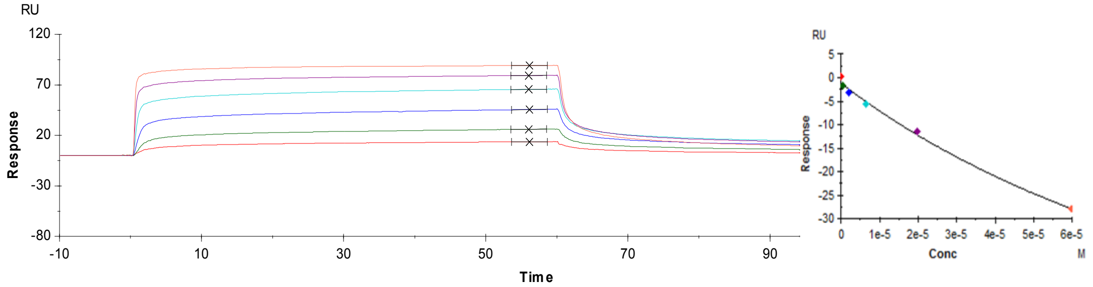


R96L

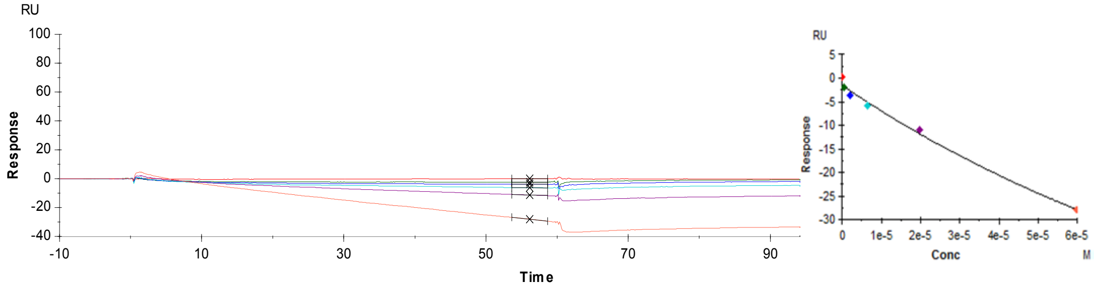


R96Q

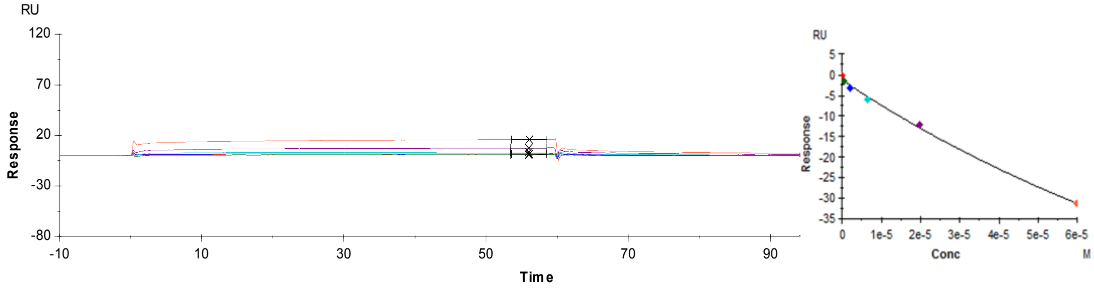


L106V

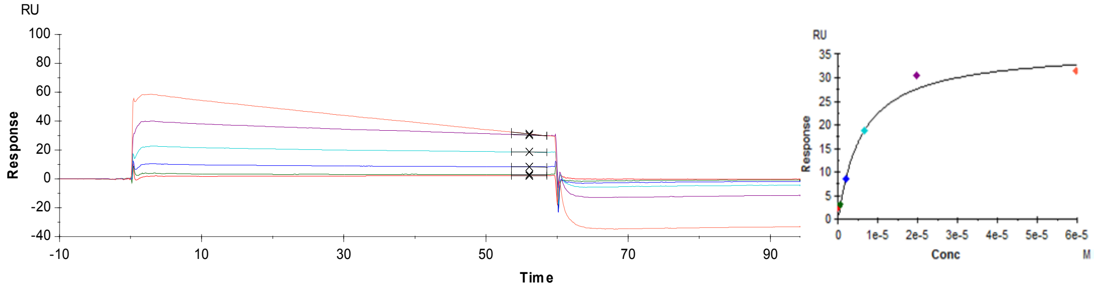


C133Y

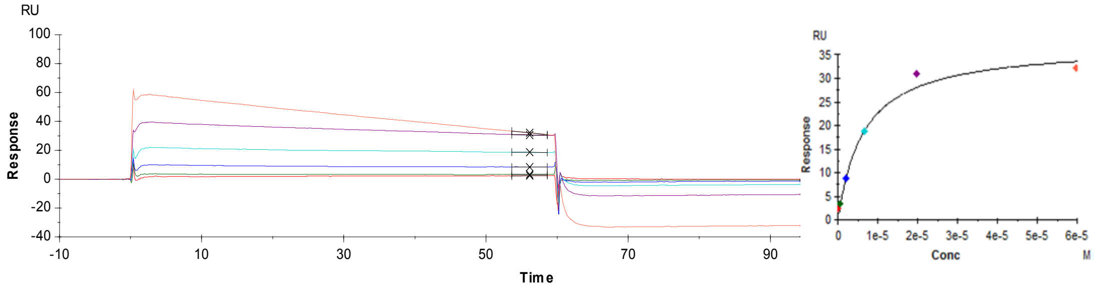


**Figure S13. The $K_D$ measurement of EpoR_pY426 peptide against SNP mutants by SPR**

Peptides were measured at concentration of 0.08, 0.25, 0.7, 2.2, 6.7 20 and 60 μM.

WT

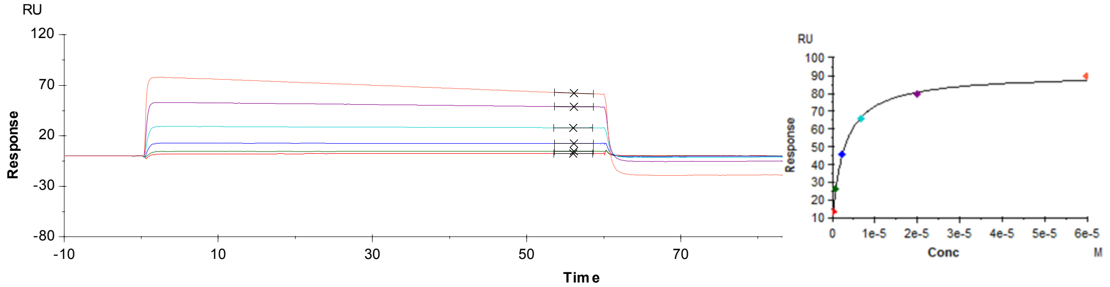


N94D

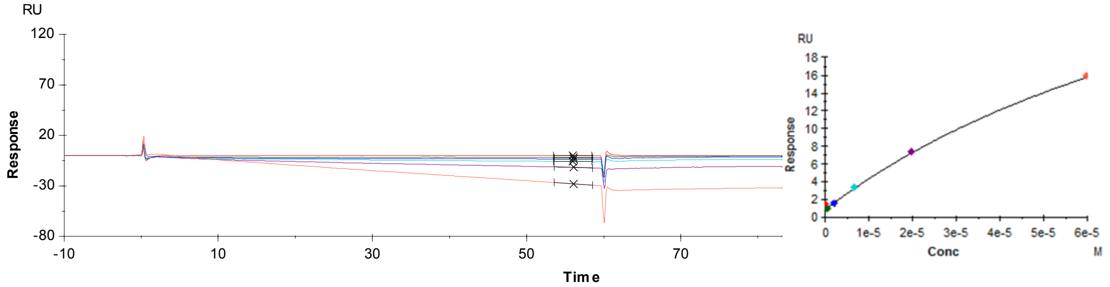


R96L

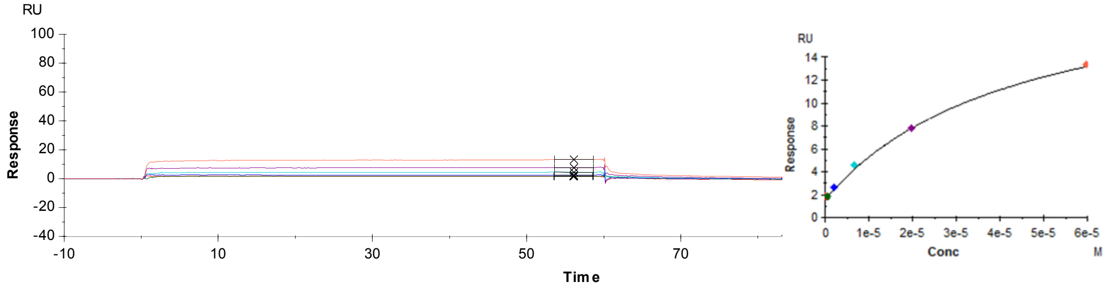


R96Q

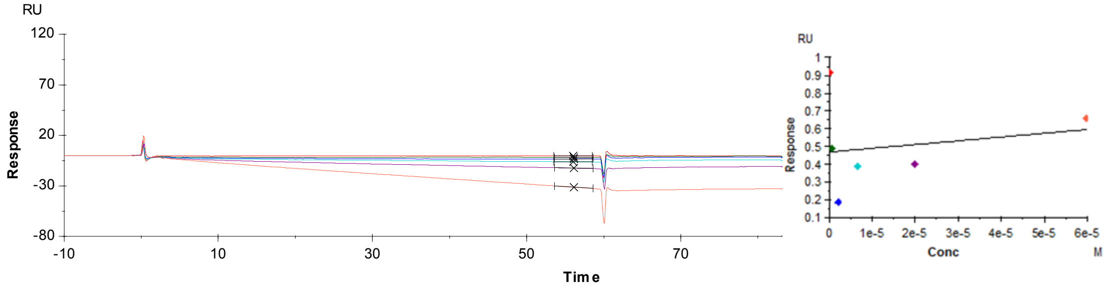


L106V

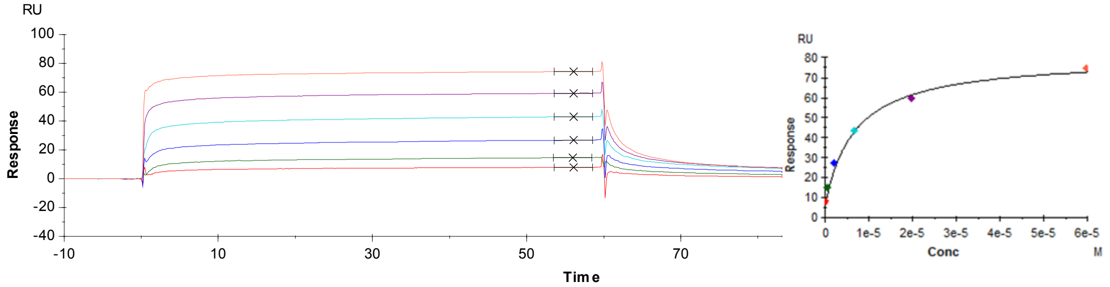


C133Y

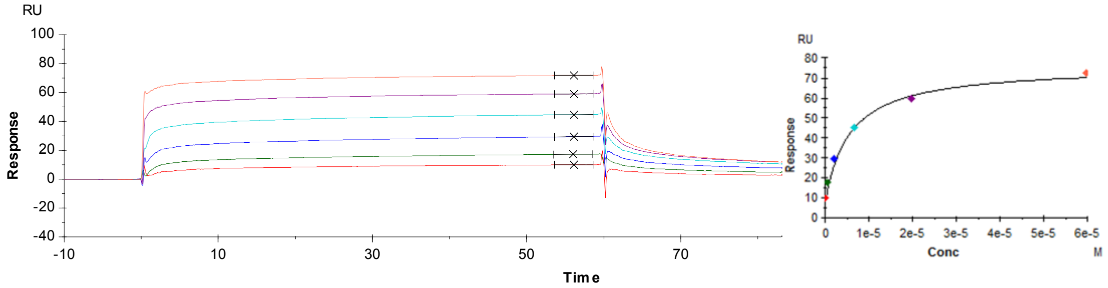


**Figure S14. The $K_D$ measurement of GHR_pY487 peptide against SNP mutants by SPR**

Peptides were measured at concentration of 0.08, 0.25, 0.7, 2.2, 6.7 20 and 60 μM.
